# The Impact of Model Assumptions in Interpreting Cell Kinetic Studies

**DOI:** 10.1101/2024.03.17.584905

**Authors:** Ada Yan, Ildar Sadreev, Jonas Mackerodt, Yan Zhang, Derek Macallan, Robert Busch, Becca Asquith

**Affiliations:** Department of Infectious Disease, Imperial College London, London, UK; Institute for Infection and Immunity, St George’s, University of London, London, UK; Department of Life Sciences, University of Roehampton, London, UK

## Abstract

Stable isotope labelling is one of the best available methods for quantifying cell dynamics in vivo, particularly in humans where the absence of toxicity makes it preferable over other techniques such as CFSE or BrdU. Interpretation of stable isotope labelling data necessitates simplifying assumptions. Here we investigate the impact of three of the most commonly used simplifying assumptions (that the cell population of interest is closed, that the population of interest is kinetically homogeneous, and that the population is spatially homogeneous) and suggest pragmatic ways in which the resulting errors can be reduced.

## Introduction

The quantification of cell dynamics is a key component of our understanding of human physiology in health and disease. Stable isotope labelling is considered the gold standard for the measurement of cell dynamics in humans *in vivo*. Stable isotope labelling has been used to investigate the dynamics of a wide range of cells, particularly haematopoietic cells, including neutrophils, monocytes and lymphocytes [1–6] and has led to fundamental discoveries in both basic cell biology [7–11] and in the pathology of diseases including HIV-1, chronic lymphocytic leukemia and type 1 diabetes [12–15].

In a typical stable isotope labelling experiment, volunteers will be given a stable isotope (usually deuterium in the form of deuterated water or deuterated glucose) for a fixed period of time: the labelling period. Blood will be drawn at multiple timepoints, during the labelling and delabelling phase, the cells of interest will be sorted by flow cytometry, DNA extracted and the fraction of deuterium-labelled nucleotides quantified by gas chromatography/ mass spectrometry. Available deuterium is incorporated when DNA is synthesised and lost when labelled cells disappear (die, change phenotype or exit the sampled compartment long-term); thus the fraction of labelled nucleotides over time contains information about cell proliferation and cell disappearance. To extract this information mathematical models are constructed and fitted to the data [16].

Mathematical models for interpreting stable isotope labelling data are necessarily simple. Over-complicated models would result in unidentifiable parameters and would not be fit for the purpose of parameter estimation. Whilst essential, these model simplifications do, potentially, impact on parameter estimates. Here we explore the impact of three widely used simplifying assumptions and suggest pragmatic ways in which the resulting errors in parameter estimates can be reduced.

The first assumption we investigate is that the cell population of interest is closed i.e. there are no upstream or downstream compartments feeding into or out of the target compartment of interest. The second assumption is that the population of interest is kinetically homogeneous i.e. all cells have the same turnover rate and the third is that the population is spatially homogeneous i.e. a sample of a population from the blood would yield the same kinetics as a sample of the same population from a spatially-distinct tissue.

## Results

### Upstream and downstream compartments

Cell populations are not isolated, closed systems: cells enter a target population as they mature and change phenotype (to the phenotype of the target population) and leave the population as they die, exit the sampled compartment long-term or change phenotype. Labelling data will often only be collected for the target cell population of interest and not for compartments upstream and downstream of the target cell compartment. There are multiple reasons why this may be the case: sometimes the lineage topology is unknown and so the identity of the upstream and downstream compartments is unknown, sometimes it is because these compartments are difficult to access in humans (e.g. maturation occurs in the bone marrow or thymus) and sometimes because blood volume is limited and so populations other than the target population cannot be studied. We have previously shown that having an upstream compartment can affect the fraction of measured label, and potentially lead to incorrect estimation of the kinetic parameters [1, 17]. Having a downstream compartment does not affect parameter estimation, provided that the disappearance rate is interpreted as the loss of cells due to all causes including exit to the downstream compartment.

Here, we investigate the conditions under which the kinetic parameters of the target cell compartment can be estimated accurately despite assuming that the target compartment is closed when model fitting. We also investigate whether fitting a model with an upstream compartment yields more accurate and/or precise estimates of the kinetic parameters compared to fitting a single-compartment model in the event that data from the upstream compartment is absent.

We consider a scenario (Figure 1) in which the target population of interest, *E*, is sampled but its upstream, precursor compartment, *C*, is not. Upstream, precursor cells *C* proliferate at rate *p*_*C*_, die at rate *d*_*C*_, and differentiate at rate Δ. Differentiation is allowed, but not required, to be linked to clonal expansion. Each precursor cell divides *k* times before differentiation (the case *k*=0 corresponds to differentiation in the absence of division). Clonal expansion is assumed to occur outside the sampled compartment (typically blood). Target population cells, *E*, proliferate at rate *p*_*E*_ and disappear (die or differentiate) at rate *d*_*E*_. The number of cells in the two populations is assumed to be independently at equilibrium. Then the dynamics of the two cell populations follow the equations

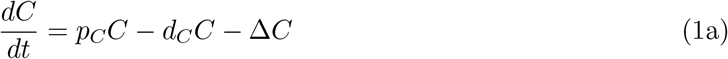

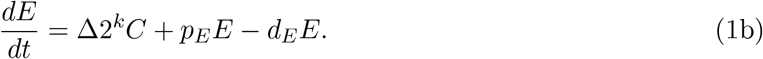

**Figure 1.**
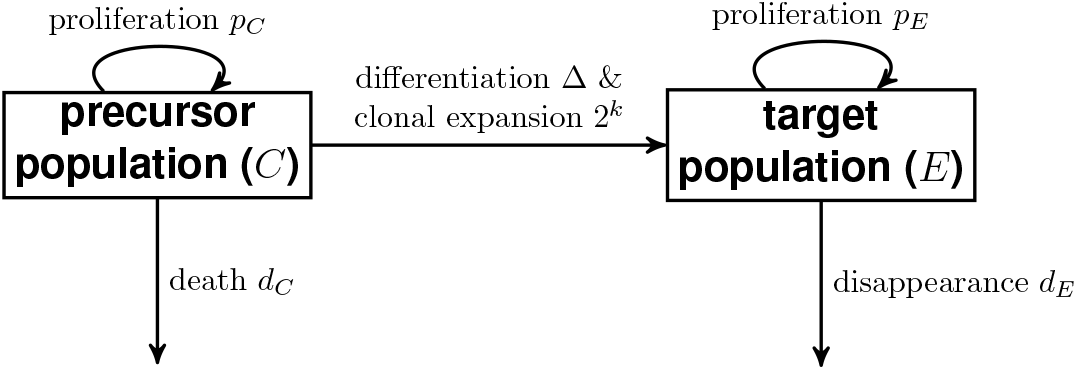
Compartmental diagram for model with precursor (upstream) and target populations.

Upon stable isotope labelling (discussed here without loss of generality for labelling with deuterated water), then the fraction of label in each compartment (*F*_*C*_ and *F*_*E*_ respectively) is

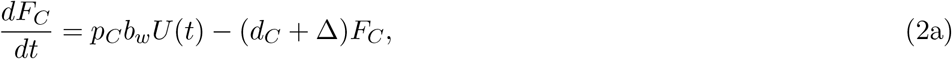

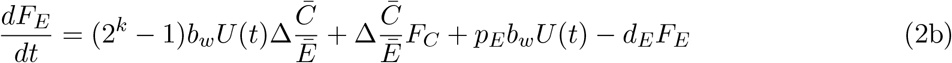

where 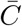 and *Ē* are the equilibrium population sizes for *C* and *E, b*_*w*_ is the normalisation factor for water [18] and *U* (*t*) is the fraction of label in body water.

We describe the fraction of deuterium in body water using a simple empirical curve

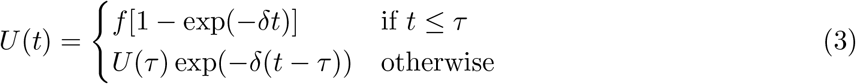

were, *f* is the plateau enrichment (expected to be the fraction of D_2_O in the daily intake of water) and *δ* the rate at which the plateau is approach (expected to be equivalent to the turnover rate of body water) and *τ* is the length of the labelling phase.

If each compartment (*C, E*) is kinetically homogeneous – that is, all cells in the compartment have the same turnover rate – then in Eqs. 1 and Eqs. 2, equilibrium constraints require *p*_*C*_ = *d*_*C*_ + Δ and 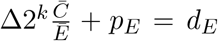. Kinetic heterogeneity in each compartment, where the precursor and/or target populations are composed of subpopulations with different turnover rates, can be approximated by replacing *d*_*C*_ and *d*_*E*_ in Eq. 2 with 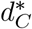 and 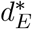 respectively where 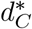 and 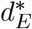 are the disappearance rates of the labelled cells in compartments *C* and *E* respectively, and we expect 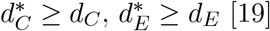.

Focusing on the target compartment (our cell population of interest), we describe its kinetics in terms of three descriptors:

i. its proliferation rate, defined to be *p*_*E*_
ii. its turnover rate, defined to be the total rate of inflow into the target compartment, 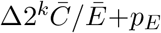, which is equal to the rate of outflow, *d*_*E*_, at steady state
iii. its rate of production by division defined to be 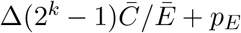.

### Derivation of relationship between estimated and actual proliferation rate when fitting a one-compartment model to the target data

When we only have measurements of *F*_*E*_ (i.e. label in the target population), one commonly used option is to neglect the existence of the precursor population and to fit a one-compartment model:

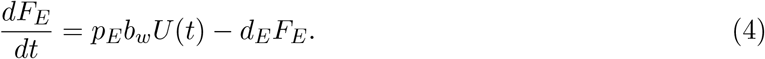

Let 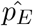 and 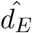 be the estimated value of the proliferation and disappearance rates obtained by fitting the one compartment model (Eq. 4) to data generated using the two compartment model (Eq. 2).

If 2^*k*^ *≫* 1, we can show that the turnover rate of the target population is very similar to its rate of production rate division, and that 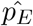 is a good approximation for both of these (see Supplementary Material). However, in this case the proliferation rate is much less than the turnover rate and so 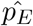 overestimates the proliferation rate. Also, the estimated disappearance rate of labelled cells, 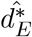, is a good approximation of the true disappearance rate of labelled cells from the target compartment, 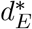.

If *k* = 0, then the proliferation rate is equal to the production rate by division, and 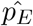 is a good approximation for both of these. However, the turnover rate is underestimated by 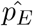, and the disappearance rate of labelled cells is not accurately captured by 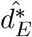.

### Confirming that the production by division is estimated accurately using numerical simulation

We sought to confirm these analytical approximations numerically. We investigated the accuracy of parameter estimation obtained by fitting a one-compartment model to two-compartment data (Methods), with kinetic heterogeneity in both the simulated data and fitted model.

Figures 2 (A-C) show the data and model fit for an example set of parameters with *k* = 10 (Supplementary Table 1). The left panel shows the simulated target population data (points) and the maximum likelihood fit (black line) when fitting Eq. 4. The one-compartment model (Eq. 4) fits the data generated by the two-compartment model (Eq. 2) very well (Figure 2A) and gives no hint of the model misspecification. However, the middle panel (Figure 2B) shows that the true proliferation rate (yellow dot-dashed line) is overestimated by 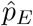, as the marginal posterior distribution (grey histogram) lies completely right of the true value. On the other hand, the estimated proliferation rate is a good estimate of the true turnover rate (dashed blue line) and the production rate by division (solid red line). The right panel (Figure 2C) shows that the estimated disappearance rate is also accurate (posterior in grey overlaps the generating value in blue).

**Table 1.** The bounds of the distribution from which transformed parameters are sampled. All rates

| Parameter | Bounds |
| --- | --- |
| $\log_{10} p_C$ | $[-4, -2]$ |
| $\log_{10} p_E$ | $[-4, -2]$ |
| $\log_{10}(\Delta/p_C)$ | $[-2, 0]$ |
| $k$ | $[0, 20]$ |
| $\log_{10}(d_C^*/(p_C - \Delta))$ | $[0, 2]$ |
| $(\log_{10} d_E - \log_{10} p_E)/(\log_{10} \delta - \log_{10} p_E)$ | $[0, 1]$ |
| $(\log_{10} d_E^* - \log_{10} p_E)/(\log_{10} \delta - \log_{10} p_E)$ | $[0, 1]$ |

**Figure 2.**
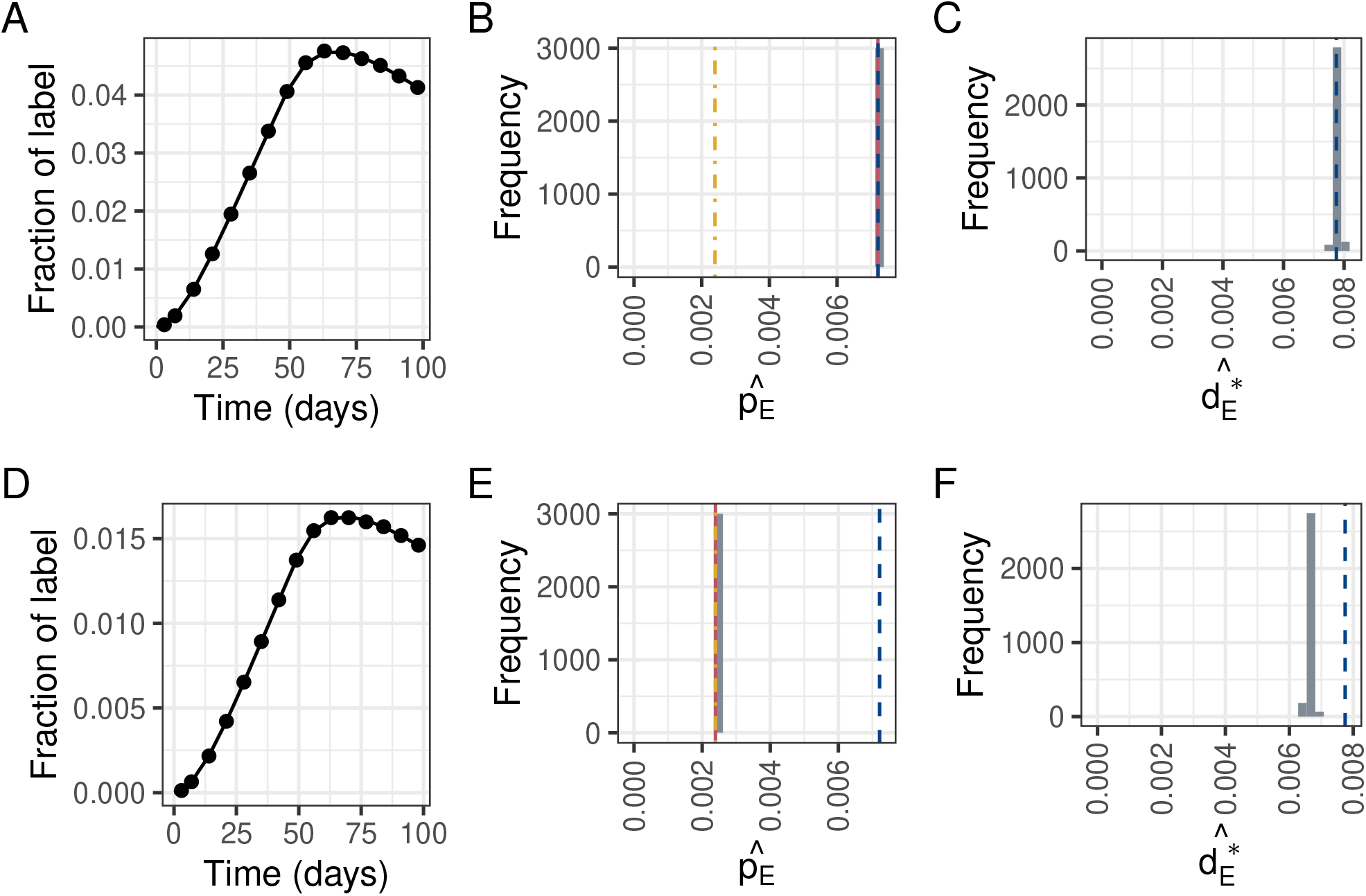
(Left. A and D) Points: Fraction of label in the target compartment generated using Eq. 2 for an illustrative set of parameters (given in Supplementary Table 1) in two scenarios: one where substantial expansion accompanies differentiation, *k* = 10 (upper row), and one where differentiation occurs without any division, *k* = 0, (bottom row). Black line: the maximum likelihood fit when fitting Eq. 4 to the data. (Middle. B and E) Marginal posterior distribution for 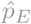 when fitting Eq. 4 to the data. The dot-dashed yellow, dashed blue, and solid red lines indicate respectively the true rates of proliferation, turnover, and production by division used to simulate the data. (Right. C and F) Marginal posterior distribution for 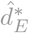 when fitting Eq. 4 to the data. The dashed blue line indicates the true disappearance rate of labelled cells used to simulate the data. Top row was simulated with *k* = 10, and bottom row was simulated with *k* = 0.

Figures 2 (D-F) show the results for the case *k* = 0. In this case 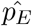 is a good estimate of the proliferation rate and the production rate by division, but underestimates the turnover rate; 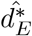 is not a good estimate of the disappearance rate of labelled cells.

Figure 3 shows that the results in Fig. 2 generalise to a wide range of parameter values. The top row shows results for 1 *≤ k ≤* 20. As expected the estimated rate 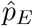 is a poor estimate of the true proliferation rate of the target compartment *p*_*E*_, consistently overestimating it (so it can be considered as a reliable upper bound). The estimated rate 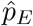 is, however, a good estimate of the turnover rate of the target compartment 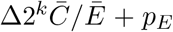 and of the rate of production of target cells by division Δ(2^*k*^ *−* 1)*C/E* + *p*_*E*_ (top row, Figure 3). Also, the estimated rate 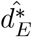 is a good estimate of the disappearance rate of labelled cells from the target compartment.

**Figure 3.**
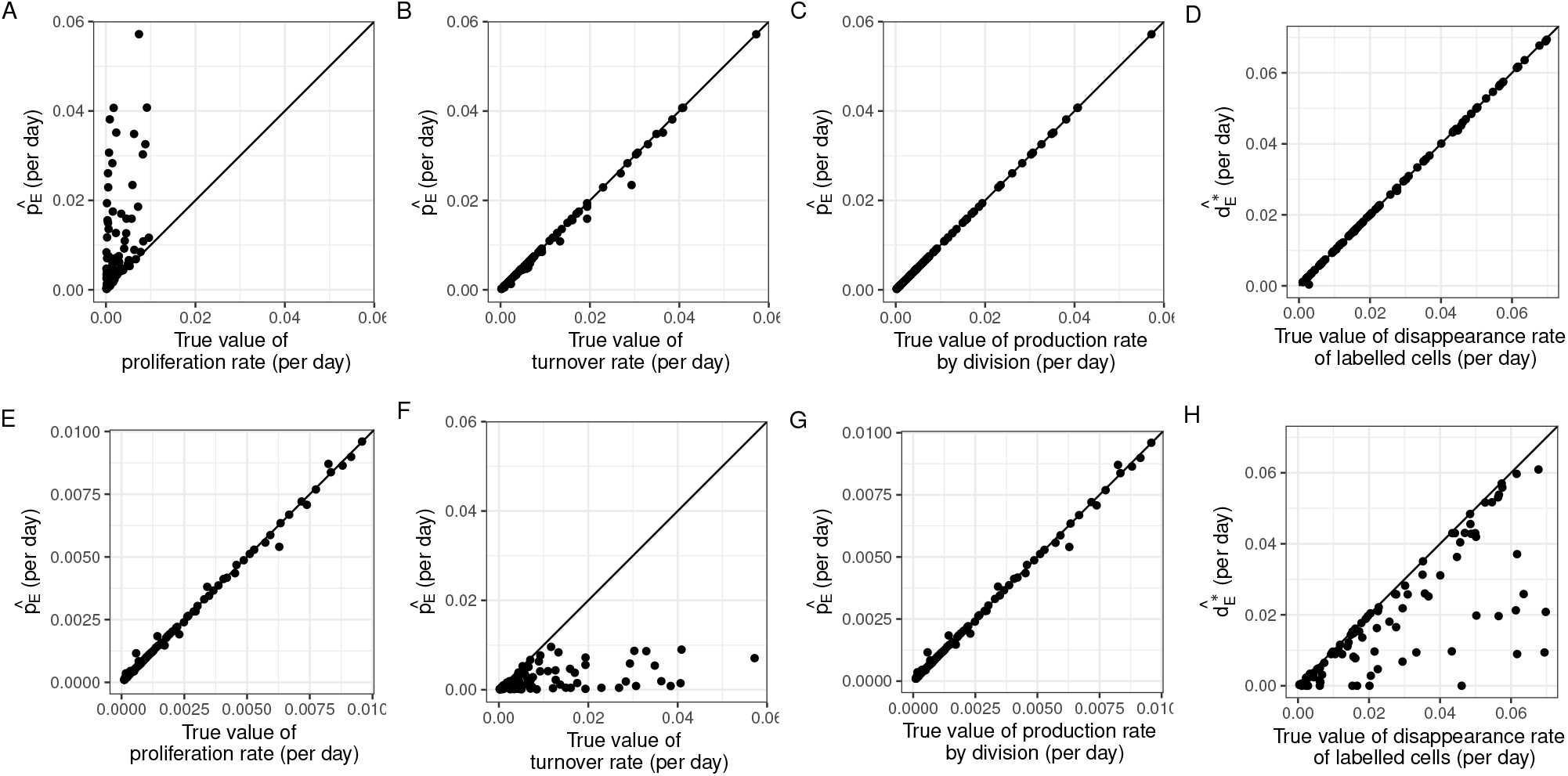
Estimated 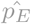 (all except rightmost column) and 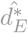 (right column) compared to (from left to right) the rates of proliferation, turnover, production by division and the disappearance rate of labelled cells in the target compartment. Top: for the case when *k* (the number of rounds of cell division associated with differentiation) varies in the range [1,20]; bottom: when *k* is constrained to zero.

The bottom row shows results for *k* = 0. As expected, turnover is now underestimated by 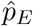 but the proliferation rate and the rate of production by cell division are accurately captured. The disappearance rate of labelled cells was also underestimated by 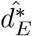.

We conclude that, when there is a significant flow of labelled cells into the target population of interest, then we (1) cannot estimate proliferation of the target population but (2) can calculate an upper bound on the proliferation rate and (3) can also estimate target cell production by division which is approximately equal to the turnover. The approximation between turnover and production by division holds provided there is division linked differentiation. The larger the value of *k* (the number of cell divisions upon differentiation) the better this approximation.

### Comparing fits using models with and without an upstream compartment

Next we investigate whether, if proliferation rather than turnover is the quantity of interest, and clonal expansion contributes significantly to label (*k ≫* 0), there is any benefit in fitting the two compartment precursor/target model (Eq. 2) despite an absence of data from the upstream precursor compartment. That is, we consider the same simulated data as in the previous section, but fit the precursor/target model. Unsurprisingly, when performing Bayesian inference, the chains do not converge so we utilised a maximum likelihood approach instead. This allows us to obtain point estimates 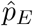 and 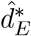, but fit diagnostics indicate that parameters are not identifiable due to colinearity and the covariance matrix is singular. Although the diagnostics indicated that the fits are unreliable, the point estimates of the proliferation rate from fitting the two compartment precursor/target model are closer to the true values than when fitting the one compartment model. We define the error to be the absolute value of the discrepancy between the true and estimated parameter values expressed as a percent of the true parameter value

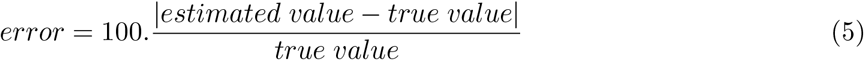

The errors in the proliferation rate estimates are significantly lower when fitting the precursor/target model than when fitting the one population model (median 201% and 83% respectively; *P* = 2.10^*−*9^, *N* = 98, Wilcoxon signed rank), Figure 4A. We reason that in an experimental setting where the identity of the precursor compartment is known but not sampled then the ratio of sizes of the precursor and target compartments can be added as a fixed parameter and that this might aid parameter estimation; we refer to this as the precursor/target model with ratio. Unexpectedly, providing the ratio of the size of the upstream to target compartments does not result in a reduction in error. If anything the errors in parameter estimates from the precursor/target model with ratio are slightly higher than the estimates from the precursor/target model without the ratio (median 109 % and 83% respectively; *P* = 0.04, Wilcoxon signed rank). However, errors still remain significantly lower than for the one compartment model, Figure 4A.

**Figure 4.**
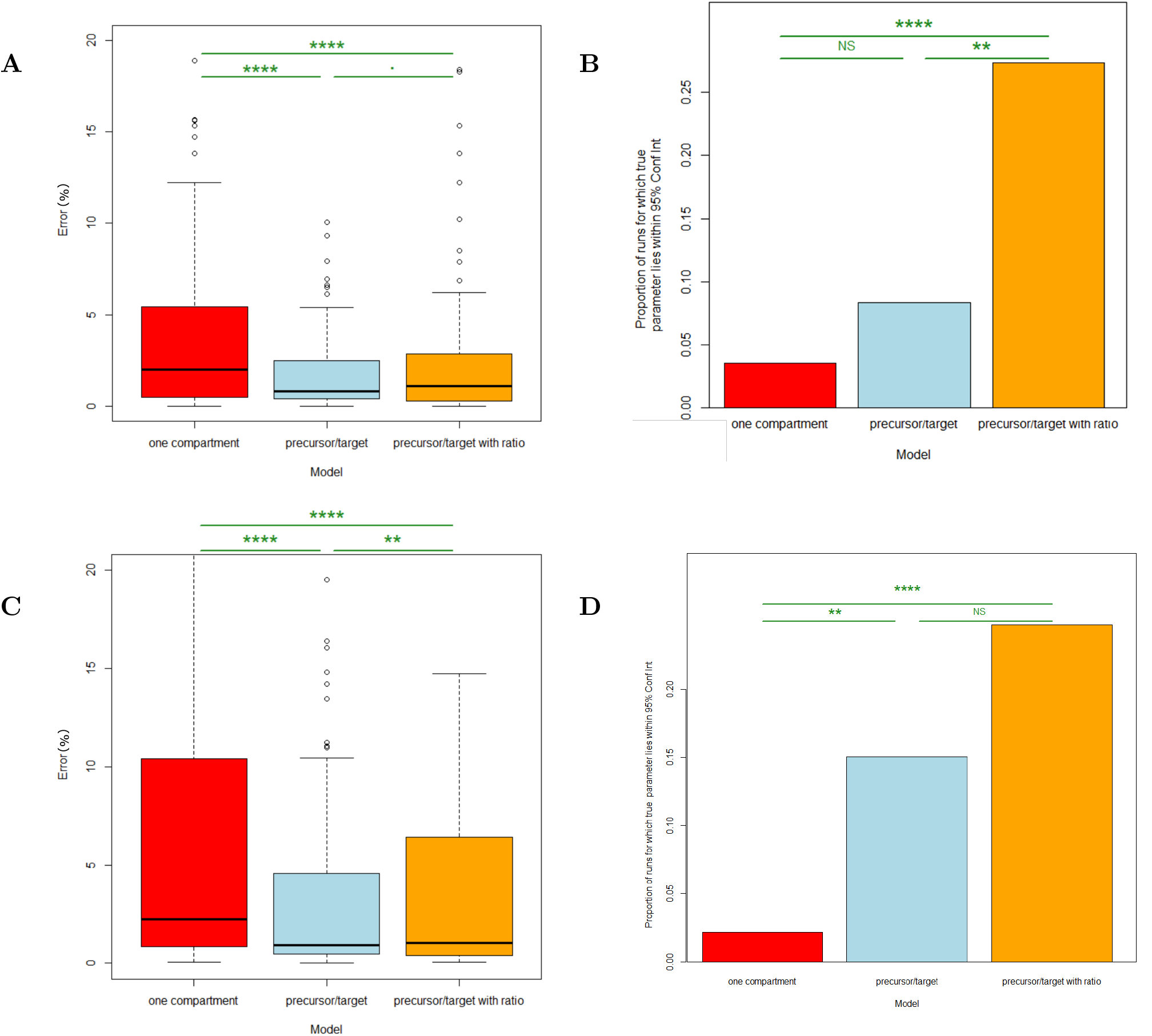
Error in estimates of the proliferation rate. **A** Discrepancy between point estimate of the proliferation rate and the true value for optimal data (expressed as a percentage of the true value) for the one compartment target model (red), precursor/target model (blue) and precursor/target with ratio model (orange). Note the y axis has been truncated to show the bulk of the data more clearly; the same graph but with no y axis truncation is given in Supplementary Figure 1. **B** 95% confidence intervals (CI) were estimated by bootstrapping the data (Methods) and the fraction of runs where the true value lay within the CI is reported. Colours as for A. The analysis in A and B was then repeated for more realistic data in **C** and **D**. Significance codes: NS *P >* 0.05, * 0.01*< P ≤* 0.05, ** 0.001*< P ≤* 0.01, ** *0.0001*< P ≤* 0.001, **** 0.00001*< P ≤* 0.0001

For each of the three models, for the 100 data sets we also estimate what proportion of times the true parameter values lay within the 95% confidence intervals (CI) of the estimated values (Figure 4B). For all three models, the proportion of runs where the true parameter value lies within the 95% CI of the estimate is low (3.6%, 8.3% and 27% for the one compartment model, precursor/target model and precursor/target with ratio model respectively). The proportion is significantly higher for the precursor/target with ratio model than for either the one compartment model (*P* = 5 *×* 10^*−*5^, Fishers exact test) or the precursor/target model (*P* = 0.003, Fishers exact test). The low proportion of runs where the estimate lies within the CI indicates that for all three models, for the data set in question, estimating the CI by bootstrapping the data leads to an underestimate of the CI (the asymptotic covariance matrix method on the other hand finds infinite intervals).

To this point we have been fitting optimal data in order to have a simpler system to aid analytical work and to focus on the effect of an upstream compartment rather than on imperfect data. There are three main respects in which our simulated data so far has been optimal: 1) noise is low 2) sampling is frequent 3) heterogeneity in the target compartment is represented by a single exponential with *d*^*∗*^ *> p* rather than explicitly with subpopulations). We therefore repeat the analysis using more realistic data by relaxing these three constraints (Methods). On calculating the errors for all three models fitted to this more realistic data we find the same pattern: i.e. the errors in the estimates of the proliferation rate obtained by fitting the precursor/target model are significantly lower than those obtained by fitting either the one compartment model or the precursor/target model with ratio (Figure 4C). Additionally, as for the optimal data, we also find that the highest proportion of runs where the true value of the parameter lies within the 95% CI of the estimated value is seen for the precursor/target model with ratio (Figure 4D). However, when changing the parameter ranges with which the simulated data are produced so that the mean proliferation of the target population is *p*_*E*_ = 0.02 (compared to a mean of *p*_*E*_ = 0.002 for the previous data sets) then relative errors produced by all models are much smaller and there are no longer any significant differences in the errors between the models (Supplementary Figure 2). We conclude that, at least for values of proliferation that are typically seen with T cells (i.e. the results shown in the main text rather than the Supplementary), fitting a precursor/target model, even when only target cell data is available produces more accurate estimates (lower percentage error) of the target cell proliferation rate.

### Kinetic heterogeneity

The second assumption we consider is the assumption of kinetic homogeneity. When estimating rates of cell turnover, a common assumption is that the population is kinetically homogeneous, i.e. all cells of the population proliferate and disappear at the same rate. However, the population may actually be composed of subpopulations which each turn over at a different rate, which is termed kinetic heterogeneity in the literature [19, 20].

Kinetic heterogeneity can be modelled explicitly. The dynamics of the subpopulations obey

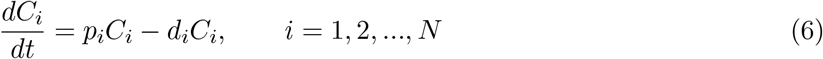

If each subpopulation is independently at equilibrium, *p*_*i*_ = *d*_*i*_, and we can define the relative size of each population as 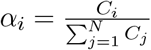. This model is illustrated for two subpopulations in Fig. 5.

**Figure 5.**
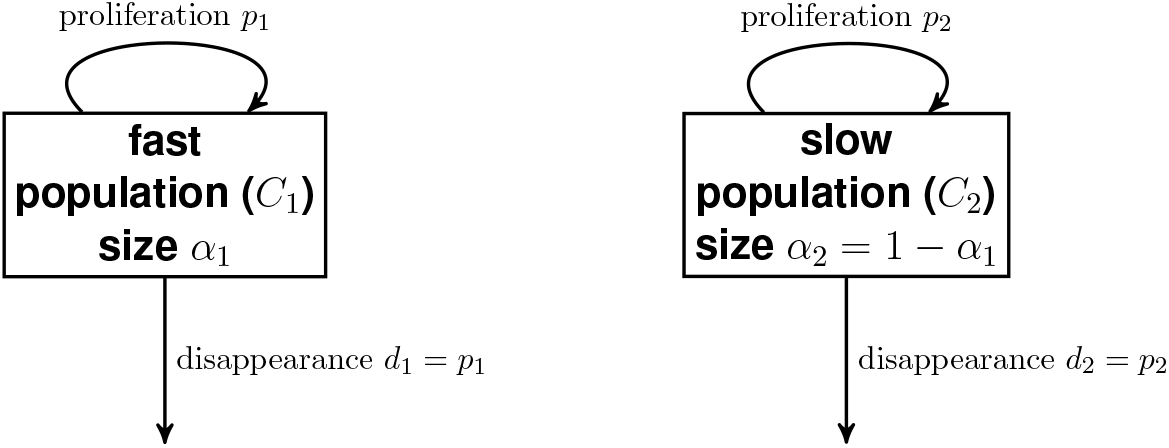
Compartmental diagram for two-compartment model with explicit kinetic heterogeneity.

The fraction of labelled cells in this explicit kinetic heterogeneity model is [20]

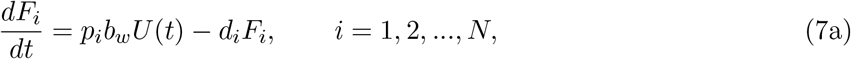

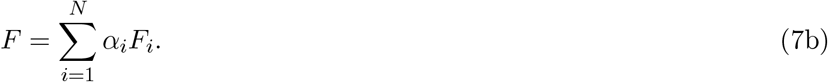

Here, *F*_*i*_ is the fraction of labelled cells in subpopulation *i* and *F* is the fraction of labelled cells in the total population.

The number of parameters scales linearly with the number of subpopulations and moreover there are strong correlations between the parameters, so some parameters may not be identifiable with limited data. A simplification can be made to yield the implicit kinetic heterogeneity model Eq. 8 [19], where *F* is the overall fraction of labelled cells, *p* is the (weighted) mean proliferation rate across subpopulations and *d*^*∗*^ is the disappearance rate of labelled cells, which is biased towards cells with the fastest turnover.

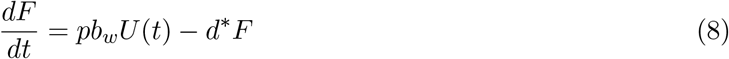

Previous studies have addressed the implications of kinetic heterogeneity on parameter estimation [6, 21–23]. However, key questions which remain unaddressed are

- How do parameter estimates obtained by fitting the implicit and explicit kinetic heterogeneity models differ?
- Can we obtain accurate parameter estimates without prior knowledge of the kinetic substructure (true number of compartments)?

### How do parameter estimates obtained by fitting the implicit and explicit kinetic heterogeneity models differ?

#### Implicit kinetic heterogeneity model parameter estimates are more precise, but potentially at the cost of accuracy

For a cell population consisting of subpopulations individually in equilibrium then the explicit kinetic heterogeneity model encapsulates the true dynamics of the system. By reducing the number of model parameters, the implicit kinetic heterogeneity model represents an approximation that potentially results in inaccurate estimates. This was shown by Westera *et al*. [23] who simulated data using a two-compartment version of the explicit kinetic heterogeneity model (i.e. *N* = 2 in Eq. (7)), then fitted explicit kinetic heterogeneity models with different numbers of compartments (*N*), as well as the implicit kinetic heterogeneity model, to the simulated data. Both the one-compartment (*N* = 1) explicit kinetic heterogeneity model and the implicit kinetic heterogeneity model underestimated the mean proliferation rate; confidence intervals from models with two or more compartments did contain the true value of the mean proliferation rate, but the uncertainty was large. Although the explicit kinetic heterogeneity model was used to generate the data and so fitting with the generating model would typically produce excellent results, the accuracy of the explicit kinetic heterogeneity model in this scenario is still surprising since it suffers from such high correlations between the parameters. We sought to explore this further.

Figure 6 shows the mean proliferation rate estimated by either (A-C) the implicit model or (D-F) a two-compartment explicit model, when fitted to data generated by a two-compartment explicit model, for a wide range of parameters (whereas Westera *et al*. used a single set of parameter values). The baseline parameters are given in Supplementary Table 2; in the panels, (left) *p*_1_, (middle) *p*_2_, (right) *α*_1_ are changed respectively. The explicit model is accurate on all counts (the 95% credible interval contains the true value), while the implicit model is only accurate for some parameter values. However, credible intervals using the explicit model are extremely wide, sometimes occupying the majority of the parameter search space (Figure 6[D-F]), whilst those from the implicit model are narrower. Hence, when using the implicit model, precision may be obtained at the cost of accuracy. Supplementary Figures 3-8 in the Supplementary Material show results for a wider range of parameter values.

**Table 2.** The parameter ranges for calculating the discrepancy between label in blood and label in lymphoid tissue. All rates are in units day^*−*1^.

| Parameter | Bounds |
| --- | --- |
| $\log_{10} p$ | $[-4, -1]$ |
| $\log_{10} f$ | $[-4, +8]$ |

**Figure 6.**
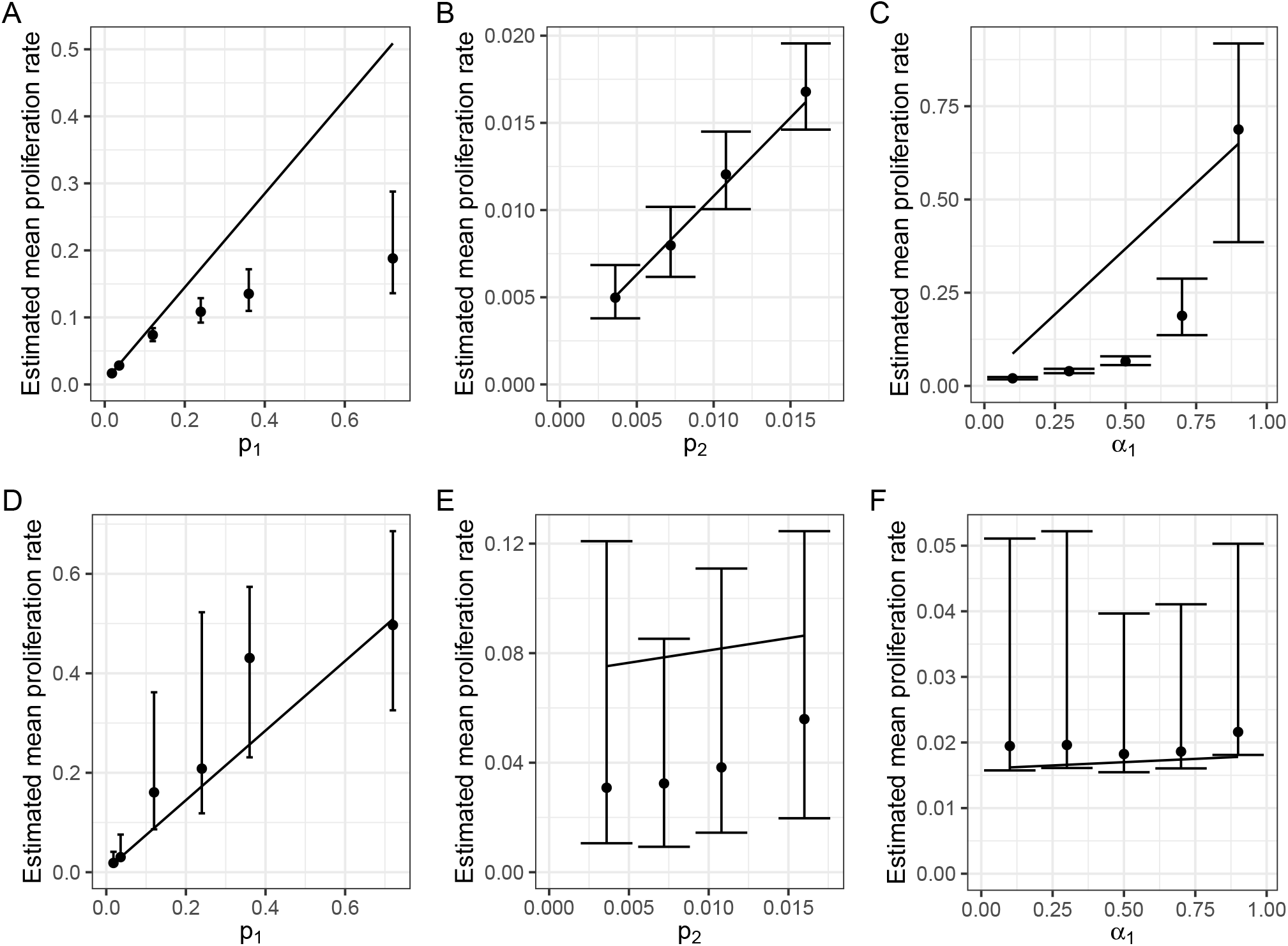
Median and 95% credible intervals for the mean proliferation rate when fitting. (A-C) the implicit kinetic heterogeneity model and (D-F) the explicit kinetic heterogeneity model with *n* = 2 to data generated using the latter model. (A, D) the value of *p*_1_ used to generate the data is changed while the remaining parameters are held constant; (B, E) *p*_2_ is changed; (C, F) *α*_1_ is changed. The diagonal lines show the true mean proliferation rate used to generate the data.

This work (Figure 6) illustrates a fundamental problem with estimating the mean proliferation rate using stable isotope labelling: that it is difficult to rule out the existence of a fast, small compartment which, while not changing the overall proportion of labelled cells much, does drive up the mean proliferation rate. On one hand, this means that confidence intervals for the mean proliferation rate for explicit kinetic heterogeneity models are wide; on the other hand, it makes the implicit kinetic heterogenity model prone to large errors, as it is inaccurate when there is a fast, small compartment.

#### Explicit kinetic heterogeneity model parameter estimates are very sensitive to the choice of prior distributions

We observed (Figure 6[D-F]) that the 95% CIs obtained by fitting the explicit kinetic heterogeneity models could approach that of the prior distribution, so we explored this further. We first simulated data from a two-compartment explicit heterogeneity model using the same parameter values as Westera et al. [23] (Methods). In line with this previous study, perfect labelling was assumed, i.e. *U* (*t*) = 1 during the labelling period and *U* (*t*) = 0 otherwise. We then used a Bayesian approach to infer the posterior distribution for the mean proliferation rate, either using the implicit kinetic heterogeneity model, or explicit models with 1-6 compartments. For consistency with Westera *et al*. we use a uniform prior distribution across [0, 1] day^*−*1^ for *p*_*i*_ (explicit model), *p* and *d*^*∗*^ (implicit model). The results are shown in Fig. 7A and are in excellent agreement with Westera *et al*. with the explicit models showing good accuracy albeit with very wide credible intervals. However, upon widening the prior from [0, 1] to [0, 10] we found that the accuracy of the explicit kinetic heterogeneity models broke down resulting in very inaccurate estimates and extremely wide credible intervals (Fig. 7B). This indicates that the estimates of mean proliferation rate obtained using the explicit heterogeneity models are heavily dependent on prior assumptions rather than the data, consistent with the high correlations between model parameters. Given that CFSE dilution studies of T cells, NK cells and B cells, both *in vitro* and *in vivo*, routinely report small subpopulations of cells undergoing large numbers of divisions (e.g. *≥* 6 divisions in 3 days, where 6 divisions represents the limit of detection in this case) it is hard to justify a conservative prior [24–26]. This dependence on the prior is not seen for the implicit kinetic heterogeneity model which yields an estimate of the proliferation rate that is the same as that seen with the narrow prior.

**Figure 7.**
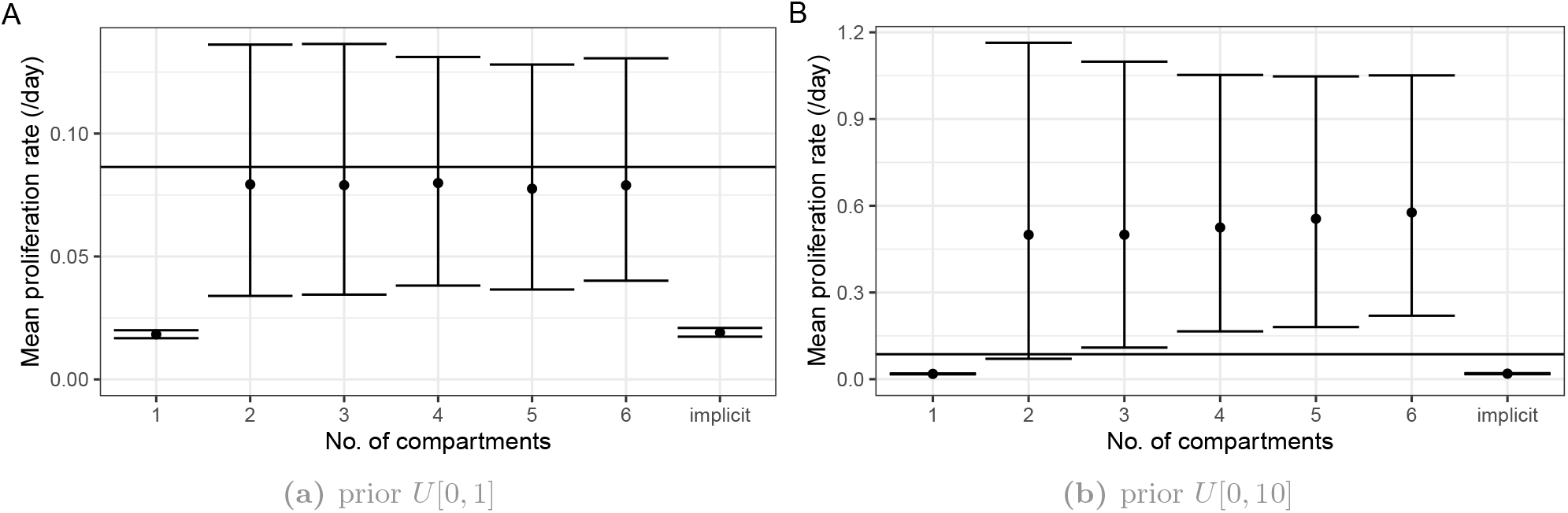
Estimated mean proliferation rates when fitting models with different priors to data generated using a two-compartment explicit heterogeneity model. Error bars indicate 95% credible intervals. Along the *x*-axis, ‘implicit’ is the fit using the implicit kinetic heterogeneity model, while the others are using explicit models with the number of compartments indicated. (A) uses a uniform prior distribution across [0, 1] day^*−*1^ for *p*_*i*_ (explicit model), *p* and *d*^*∗*^ (implicit model), and (B) uses a uniform prior distribution across [0, 10] day^*−*1^ for these parameters. The horizontal line indicates the value used to simulate the data.

#### Insights can be drawn by fitting the implicit kinetic heterogeneity model to the data

Next we looked at what insights can be drawn by fitting the implicit kinetic heterogeneity model to the data. First, whether the estimated ratio *d*^*∗*^*/p* exceeds 1 can indicate whether kinetic heterogeneity is present. However, the power of this method to detect heterogeneity heavily depends on the true parameter values, as shown in Fig. 9 in the Supplementary Material.

Second, the mean proliferation rate estimated by the implicit model serves as a lower bound for the true mean proliferation rate, and an upper bound for the proliferation rate of the slowest compartment. Figure 6A-C and Supplementary Figures 3-5 show that the mean proliferation rate estimated by the implicit model serves as a lower bound for the true mean proliferation rate. Figure 8 shows the estimated mean proliferation rate using the implicit model (y-axis) versus the value of *p*_2_ used to generate the data (labels above the plots, also shown as the horizontal line), for different values of *p*_1_ (x-axis) and *α*_1_ (in each row). All estimates of the mean proliferation rate lie above the value of *p*_2_ used to generate the data, indicating these estimates are an upper bound of the proliferation of the slowest compartment.

**Figure 8.**
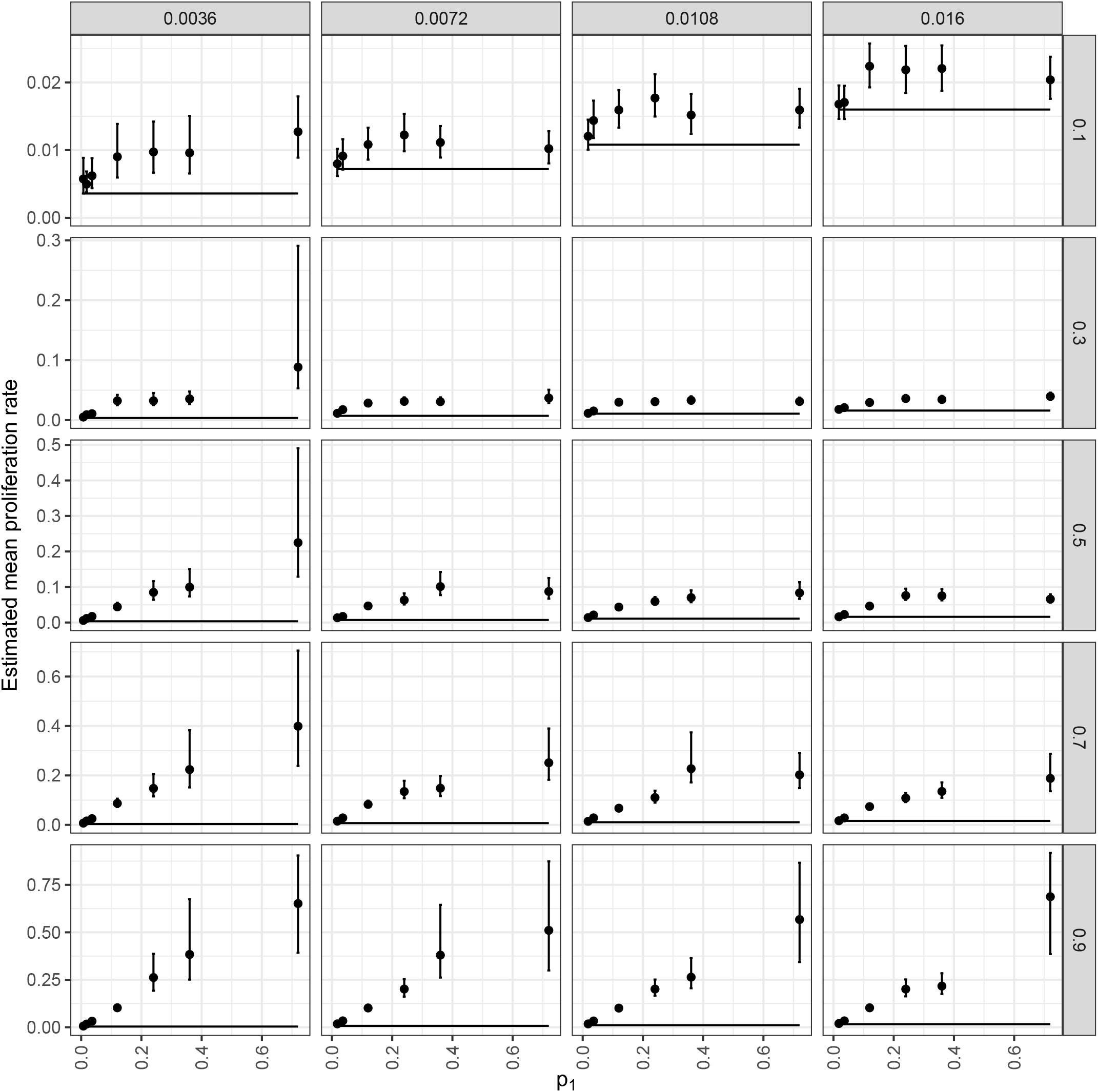
The mean proliferation rate estimated by the implicit model serves as an upper bound for the proliferation rate of the slowest compartment. Estimated mean proliferation rate using the implicit model (y-axis) vs value of *p*_2_ used to generate the data (horizontal lines and labels above the plots), for different values of *p*_1_ (x-axis) and *α*_1_ (different rows). Dots and error bars indicate median and 95% credible intervals respectively.

**Figure 9.**
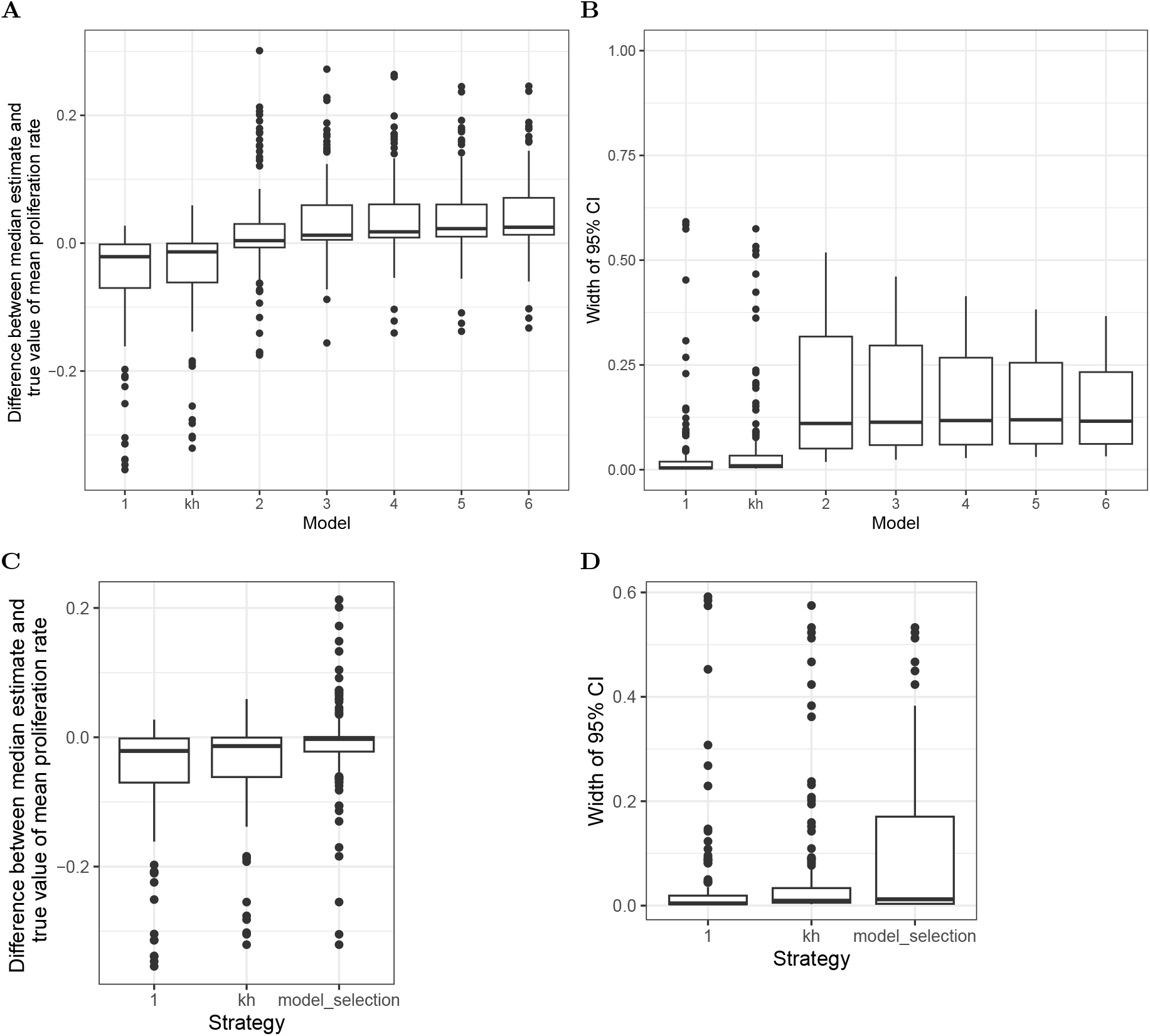
(A) The error (difference between the estimated proliferation rate and the true, generating, proliferation rate) for each model used to fit the simulated data generated with the n=2 explicit heterogenity model. (B) The width of the 95% CI for each model (C) The error in the proliferation rate, using either of the simplest two models or using model selection. (D) The width of the 95% CI of the proliferation rate, estimated using either of the simplest two models or using model selection. ‘kh’ denotes the implicit kinetic heterogeneity model whilst the numbers indicate the number of compartments in the explicit model (so “2” is the generating model).

Finally, when comparing turnover kinetics between two data sets – for example, between healthy individuals and individuals with a given condition – the implicit model is able to pick up differences in the mean proliferation rate, if it is assumed that the two data sets differ in only one parameter (*p*_1_, *p*_2_ or *α*_1_). Figure 6A-C shows the mean proliferation rate estimated by the implicit model, when fitted to data generated by the explicit model, as (A) *p*_1_, (B) *p*_2_, (C) *α*_1_ is changed. We see that the mean proliferation rate estimated by the implicit model increases as each of these parameters is changed individually, which also increases the mean proliferation rate of the explicit model. Even though the quantitative difference is incorrect, the estimated changes are in the correct direction. By contrast, Fig. 6D-F shows that fitting the explicit model to each data set separately does not allow us to distinguish changes, as the confidence intervals are so wide. The explicit model might be able to pick up these differences if it is fitted simultaneously to both data sets and the assumption that only one parameter value is different between the two is encoded into the model; this is the subject of future work.

### Can we obtain accurate parameter estimates without prior knowledge of the kinetic substructure (true number of compartments)?

For real data, we do not know the number of kinetically distinct subpopulations. One approach, suggested by de Boer *et al*. [22], is to increase the number of compartments in the fitted explicit kinetic heterogenity model until the goodness of fit is no longer improved. We investigate the effectiveness of this approach.

We simulate data using a two-compartment version of the explicit model, using different values of *p*_1_, *p*_2_ and *α*_1_ (Methods). We then fit either an *n*-compartment explicit kinetic heterogenity model or the implicit kinetic heterogeneity model to the data. For computational feasibility the number of compartments, *n*, is capped at 6.

Before applying the model selection strategy proposed [22], we first examine both the error and the precision (CI width) achieved when fitting each of these models to the 125 simulated data sets.

Figure 9A shows the error (difference between the estimated proliferation rate and the true, generating, proliferation rate). As expected, the generating model (explicit model with 2 compartments) is most accurate. The single-compartment version of the explicit model and the implicit kinetic heterogeneity model tend to underestimate the proliferation rate and the explicit models with *n ≥* 3 tend to overestimate the proliferation rate. Surprisingly, adding more compartments to the explicit model increased the error, even though models with more compartments are more general versions of the model from which the data was generated. This is because using stable isotope labelling data, it is difficult to rule out the existence of small populations with fast turnover, which increase the mean proliferation rate but hardly increase the amount of label due to their small size and label saturation. Therefore, allowing the explicit model more flexibility with additional compartments increases the estimate of the mean proliferation rate, if the same prior distribution is used for the proliferation rate for all compartments. Figure 9B shows that the width of the 95% CI is significantly narrower for the explicit one compartment model and the implicit kinetic heterogeneity models than for all the *n ≥* 2 explicit models, in line with our findings in the previous section *How do parameter estimates obtained by fitting the implicit and explicit kinetic heterogeneity models differ?*.

Next we investigate model selection strategies. We compare the effectiveness of always using the simplest model (one-compartment explicit heterogeneous); always using the implicit kinetic heterogeneity model; and model selection of the explicit kinetic heterogeneity models on the basis of good-ness of fit. Model selection is implemented by first considering the two simplest models (*n* = 1 and *n* = 2)where we define a model’s complexity by the number of fitted parameters. We use 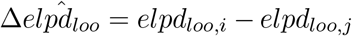, which is a metric related to the expected log pointwise predictive density, as the criteria to compare these two models (Methods). Where there is insufficient evidence to select the more complex model, the simpler model is selected; where there is evidence to select the more complex model, the next most complicated model is added to the set of models under consideration, and model comparison is repeated. This process is repeated until the most complex model within the set of models under consideration is no longer favoured statistically, leading to selection of the second most complex model in the set.

Figure 9C compares the error in the model selection estimate versus either always using the simplest model or always using the implicit kinetic heterogeneity model; whilst Figure 9D shows the width of the CI for the same three cases. Model selection has a slightly lower error in the estimates and the widest CI. However, it should be noted that, for the model selection approach, both the estimate and the CI will depend on the choice of prior (see previous section *How do parameter estimates obtained by fitting the implicit and explicit kinetic heterogeneity models differ?*). Here we have used a more conservative prior of *U* [0, 1] that is aligned with the generating data; widening the prior will result in higher errors and wider CI.

In summary, there is no model that always performs best by every metric. Individual studies will have to balance precision and accuracy and conduct an accurate assessment of prior data when choosing a model selection strategy.

### Spatial distribution of T lymphocytes

The final assumption we consider is the assumption of spatial homogeneity. Studies of T cell dynamics, at least for humans, are most often based on samples from the blood. Directly *ex vivo*, most peripheral blood T lymphocytes are in G0 *\*G1 [27, 28]. Nevertheless high levels of deuterium are typically detected in peripheral blood lymphocytes following a labelling period. This indicates that division occurs outside the blood, most likely in lymphoid tissue, and that dividing cells take up label outside the blood stream and then travel through the blood where they are sampled. Here we ask whether cell proliferation and disappearance rates, estimated from the timecourse of labelled peripheral blood cells are accurate estimates of proliferation and disappearance of cells in lymphoid tissue. That is, do labelled cells in the blood provide a window into cell dynamics in lymphoid tissue? We address this question using the following model:

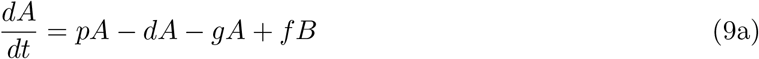

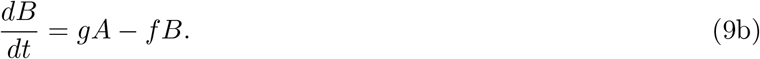

In which *A* is the number of cells (of the cell population of interest) in lymphoid tissue and *B* the number of cells of the same population in the blood. Cells in lymphoid tissue proliferate at rate *p*, disappear (die/ differentiate/ exit the circulation long-term) at rate *d*, enter the blood at rate *g* and leave the blood at rate *f*. Upon labelling, then the fraction of label in each compartment (*F*_*A*_ and *F*_*B*_ respectively) is

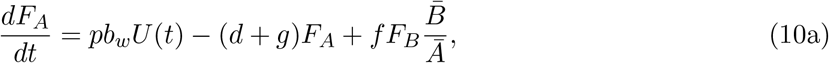

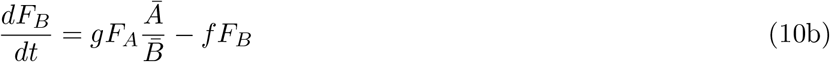

where *Ā* and 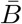 are the steady state sizes of the populations *A* and *B* (i.e. 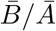 is the blood to lymph ratio for the population of interest, which is of the order of 2*/*98 for lymphocytes) and *U* (*t*) is the fraction of deuterium in body water (Equation 3). We assume that only label in the blood, *F*_*B*_ can be observed.

We simulate this system using parameters from physiological ranges (Methods) and calculate the discrepancy between the label in the lymph and the observed label in the blood 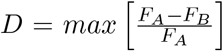 for a range of values of *f* and *p*. The maximum discrepancy is plotted in Figure 11 and a representative labelling plot for a particular set of parameters in Figure 12.

**Figure 10.**
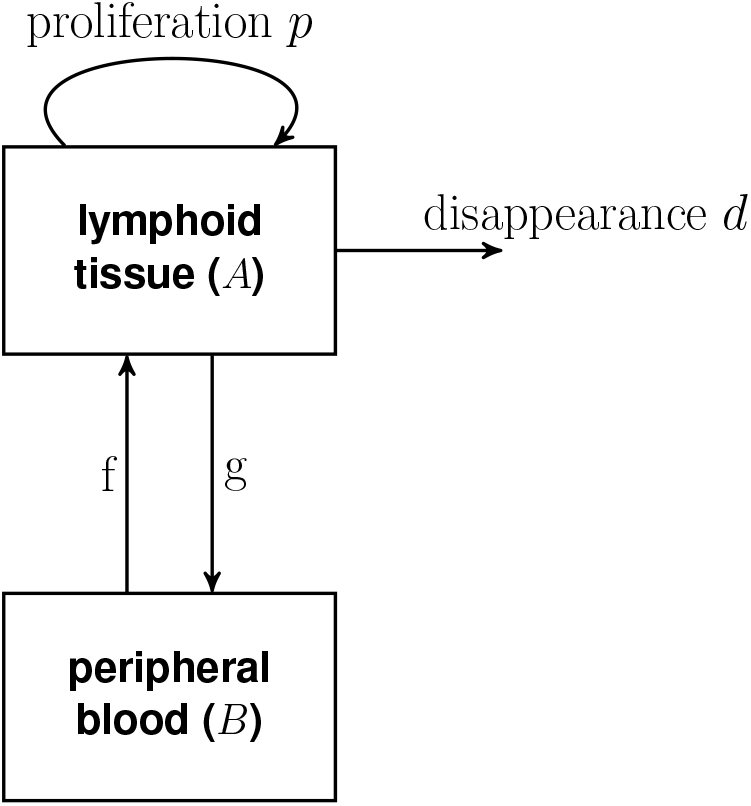
Flow diagram for model with lymphoid tissue and blood.

**Figure 11.**
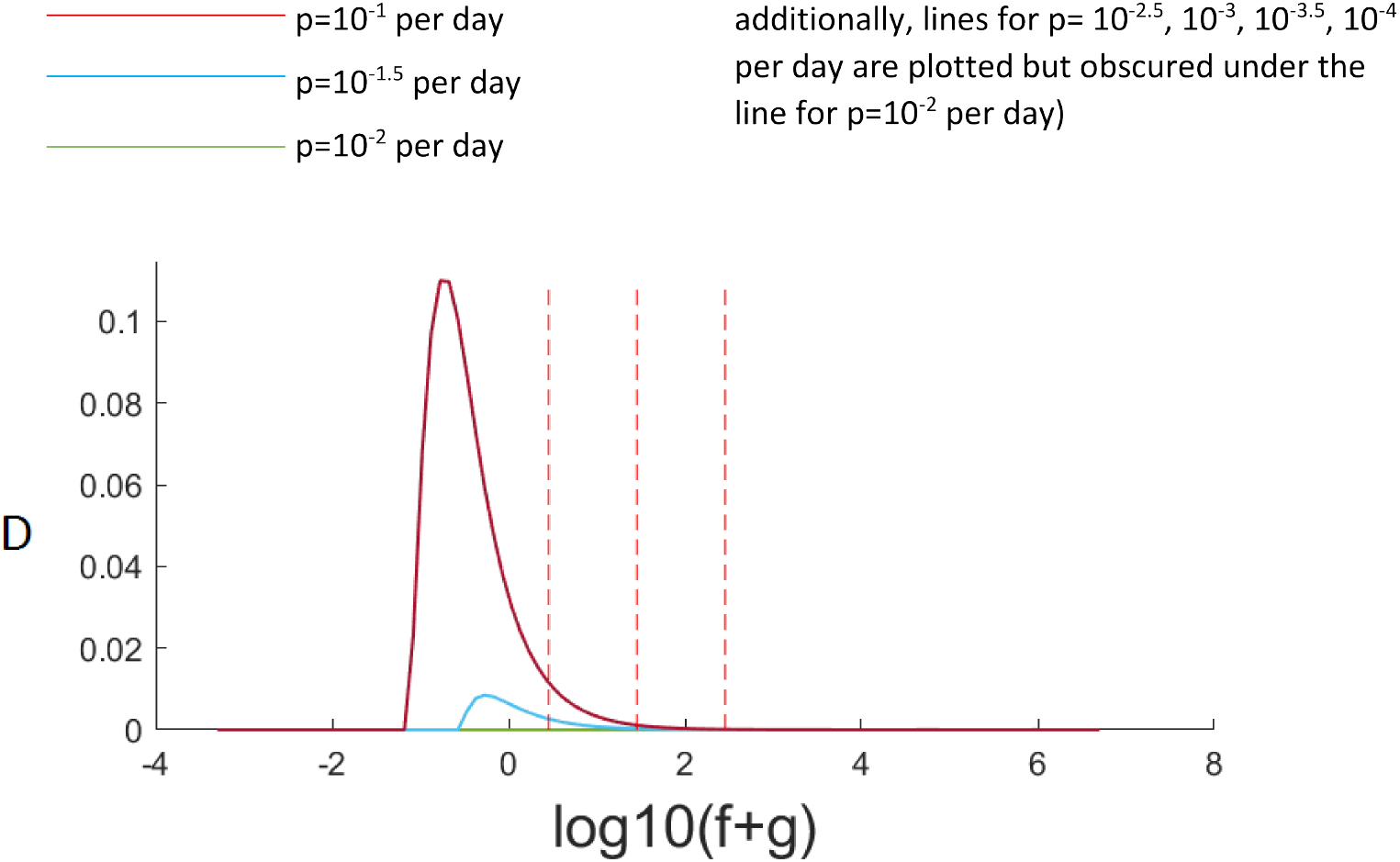
Maximum discrepancy, D, between label in blood and label in secondary lymphoid tissue for varying lymphocyte proliferation rates (different coloured lines), varying *f* + *g* along the x axis. The central vertical dashed line marks the physiological recirculation rate of f=28 per day, lines to left and right mark 10 fold lower and higher (2.8 per day and 280 per day) respectively. It can be seen that unless recirculation is unrealistically slow and the lymphocyte kinetics are very fast then the label in the blood is a very good approximation of label in the lymphoid tissue.

**Figure 12.**
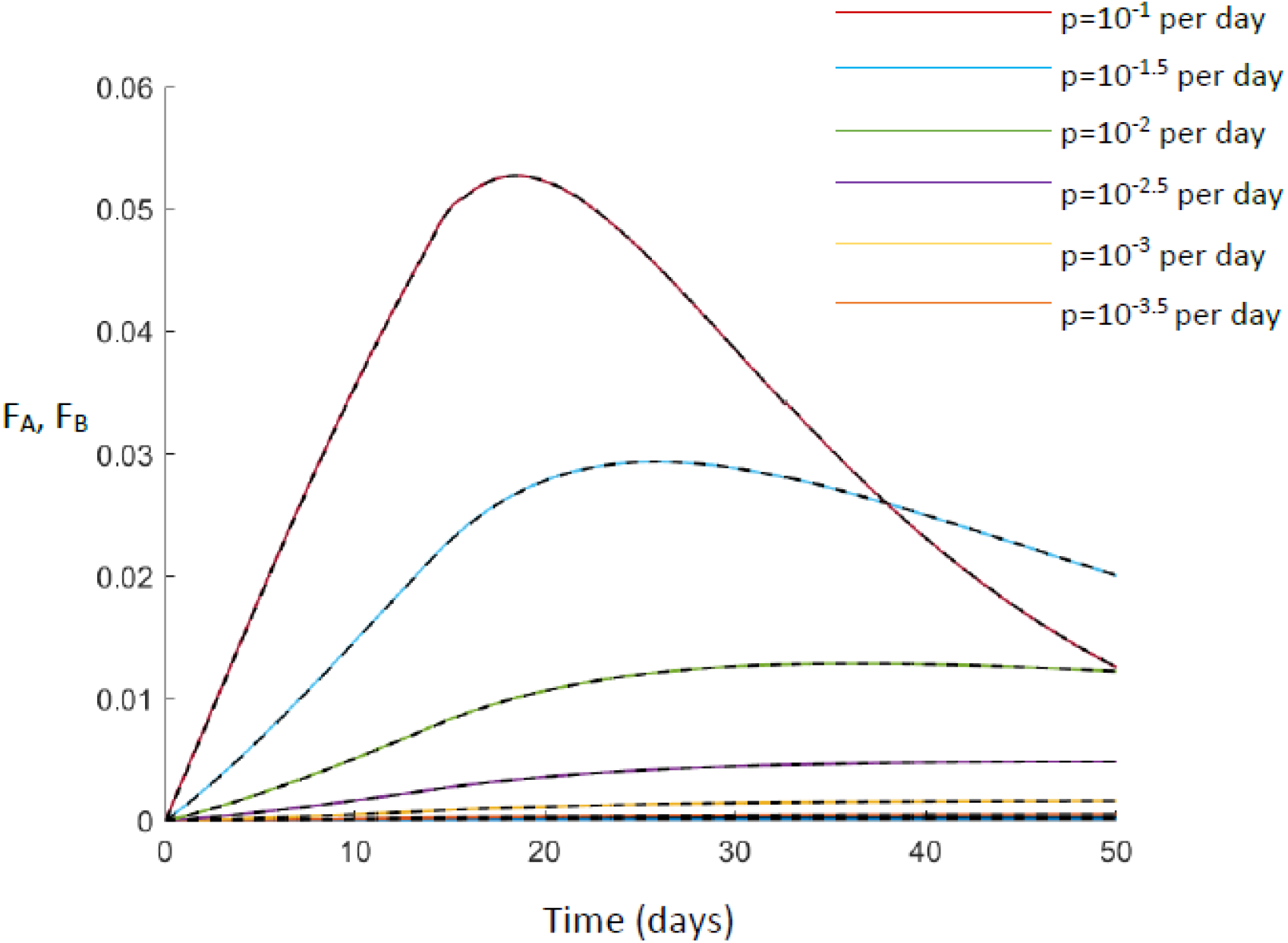
A comparison of label in lymph node (solid line, *F*_*A*_) and label in blood (dashed line, *F*_*B*_). It can be seen that, for all proliferation rates considered, they overlay. Here *f* =28 per day.

It can be seen that, for our best estimate of the lymphocyte recirculation rate (*f* =28 per day) then for all values of proliferation considered, the maximum discrepancy at any timepoint is extremely small. Even if the lymphocyte recirculation rate is ten-fold slower (*f* =0.28 per day), and the proliferation rate is high (*p*=0.1 per day) then the maximum discrepancy is still less than 2%. We conclude that, for freely recirculating T lymphocytes, then dynamics quantified from the blood will be representative of dynamics in lymphoid tissue.

## Discussion

We have investigated the impact of three widely used simplifying assumptions when modelling stable isotope labelling data. The first is the assumption that the target subpopulation being studied is closed. We considered the situation that there is an upstream precursor population flowing into the target cell population but that the upstream compartment is not sampled. This scenario can lead to large errors in the estimation of the proliferation rate of the target cell population if the precursor population is not modelled. We found that accuracy in estimating the proliferation rate depended on the number of cell divisions upon differentiation, *k*. When 2^*k*^ *≫* 1, the estimated proliferation rate provided a good estimate of the turnover rate and the rate of production by division for the target compartment, but it was only an upper bound for the true proliferation rate of the target compartment. The estimated disappearance rate of labelled cells in the target compartment was accurate. On the other hand, when *k* = 0, the estimated proliferation rate is a good estimate for the proliferation rate of the target compartment (which in this case is equal to the production rate by division), but is a lower bound for the turnover rate of the target compartment; the estimated disappearance rate of labelled cells is also a lower bound for the disappearance rate of labelled cells from the target compartment. As parameter estimates behave differently for different values of *k*, it is difficult to interpret them without independent information about *k*. However, in all cases, the rate of production by division is accurately estimated. In the case where 2^*k*^ *≫* 1, a better estimate of the proliferation rate can be obtained by fitting a precursor/target model, even when data from the precursor population is absent.

Swain *et al*. (under review) addressed the same question but restricted to kinetically homogeneous upstream/target compartments, and for the special cases of *k* = 0 and *k* = 1, using different analytic derivations. As the kinetically homogeneous *k* = 0 and *k* = 1 cases are special cases of our model (heterogeneous *k ∈* [0, 20]), the results of our two studies should agree for this case. For *k* = 0, we agree that the turnover rate may not be well estimated if the target compartment is assumed to be closed. Swain *et al*. goes further to identify the situations in which the turnover rate is well estimated regardless – when the turnover rate of the upstream compartment is fast, or when the inflow into the target compartment is small.

The second assumption we analysed is that all lymphocytes in a population have the same turnover rate. Although previous studies have used implicit or explicit models to address kinetic heterogeneity, the circumstances under which each model should be used, and their limitations, have not been investigated. If the explicit model is the generative model then, comparing the implicit and explicit models, we found that the implicit model had greater errors when estimating the mean proliferation rate, but CIs for the explicit model were driven by prior assumptions rather than the data in hand. Fitting the implicit model enables us to assess whether there is evidence of heterogeneity in the population and whether a model with kinetic heterogeneity is required at all. It is also able to identify directional differences in the mean proliferation rate between samples, where the kinetics are assumed to differ by a single parameter between samples. Furthermore, the mean proliferation rate estimated by the implicit kinetic heterogeneity model represents a lower bound of the true mean proliferation rate and an upper bound on the proliferation rate of the smallest compartment. Comparing explicit kinetic heterogeneity models with different numbers of compartments, we found that as expected, credible intervals for the mean proliferation rate from a model with more compartments were more likely to contain the true mean proliferation rate. However, if model selection criteria were used to select the most statistically supported model, more precise estimates could be obtained, although for a small proportion of parameter sets, the true value is no longer contained in the 95% credible interval. Selecting the most appropriate model thus involves balancing accuracy with precision.

The third assumption is the assumption that lymphocytes are spatially homogeneous and in particular the assumption that the fraction of labelled lymphocytes quantified in the blood is representative of the fraction of labelled lymphocytes in the secondary lymphoid tissue. We found that this assumption was reasonable for lymphocytes that recirculated between blood and lymphoid tissue at a rate of about 0.3 per day or faster. This is slow compared to the average rate of lymphocyte recirculation, so, provided a subpopulation is not recirculating much slower than average its kinetics should be reliably quantified from blood samples. If a lymphocyte population is sequestered in lymphoid tissue or only recirculating very slowly then label in the blood will be poorly representative of label in the secondary lymphoid tissue.

In summary, we find that some frequently used assumptions can have profound impact on the interpretation and estimates of parameter values. Whilst these assumptions are typically unavoidable it is important to be aware of their implications when conducting labelling studies.

## Methods

Code to reproduce results can be found at https://github.com/ada-w-yan/cellkineticmodels.

### Simulating data for precursor/target model (Section *Upstream and downstream compartments*)

Transformed versions of the upstream and downstream parameters were sampled from a multivariate uniform distribution using Latin hypercube sampling. Table 1 shows the bounds of the distribution. Bounds above the line in Table 1 are bounds that were chosen to reflect physiological estimates. They include the bounds on *p*_*C*_ and *p*_*E*_ which were chosen in line with typical estimates of T cell proliferation in healthy individuals [29]. 0.01*p*_*C*_ *≤* Δ *≤ p*_*C*_ was chosen so that the differentiation rate of the upstream population is smaller than its proliferation rate, as would be expected in homeostasis (the lower bound is imposed so there is flow between the two compartments; otherwise the problem reduces to the trivial solution of two independent compartments). The number of rounds of clonal expansion *k* was initially chosen to lie between 0 and 20. 0 represents the scenario when cells flowing from *C* to *E* are unlabelled.

Bounds below the line are necessary to reflect model constraints. 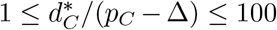 enforces the condition that 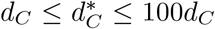 i.e. the upstream population *C* has the freedom to be kinetically heterogeneous (with the maximum heterogeneity constrained such that the death rate of labelled cells is not greater than 100 times the average death rate). 0 *≤* (log_10_ *d*_*E*_ *− log*_10_*p*_*E*_)*/*(log_10_ *δ −* log_10_ *p*_*E*_) *≤* 1 and 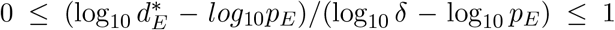 impose *p*_*E*_ *≤ d*_*E*_ *≤ δ* and 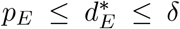 respectively. The lower bounds are necessary for the target population to remain in equilibrium and the upper bounds reflect the design of stable isotope labelling experiments which mandates that the loss of the isotope from the body (*δ*) must be more rapid than the loss of the population of interest. In addition, we only keep samples that fulfil 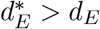. This allows the target population to be kinetically heterogeneous. The value of *δ* was fixed to 0.07 which is a realistic value from the literature [2].

Supplementary Figure 10 shows the shape of the distribution in terms of the original parameters.

We generate 100 parameter samples from the above distribution and 100 parameter samples from the version with *k* = 0. For each sample, we substituted parameter values into the analytic solution of Eq. 2 assuming a labelling period of 49 days. We take data points daily up to day 100.

#### Optimal data

Initially we simulated an optimal experimental situation with a large number of points with a small standard deviation because the aim is to determine the effect of the upstream compartment and not the effect of uncertainty in the data. We take data points daily. Each data point is lognormally distributed with parameters *μ* = log(*x*) where *x* is the true value of the fraction of label, and *σ* = 0.005.

#### Realistic Data

Although our focus was on “optimal data” we also investigated how our conclusions varied with more realistic data. The realistic data differed from the optimal data in three main respects. First, data sampling was assumed to be much sparser: we assumed measurements once every week, resulting in 15 measurements over the 100 day period of the experiment. Second, we allowed for considerably more noise: noise was normally distributed with mean zero and standard deviation 0.1*×* max value of label in that data set. Finally, we explicitly modelled heterogeneity in the target compartment. That is, we modelled two subpopulations within the target compartment (each independently in equilibrium) rather than a single population with *d*^*∗*^ *> p*. The equations to describe label in this model are as follows:

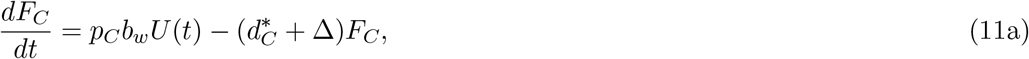

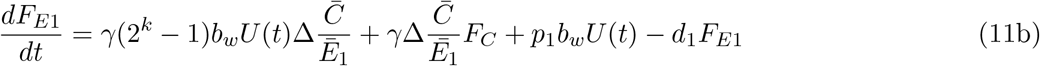

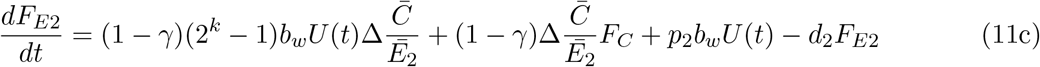

Where *F*_*C*_ is the fraction of label in the upstream compartment and *F*_*E*1_ and *F*_*E*2_ is the fraction of label in the two target subpopulations. Cells in the upstream compartment proliferate at rate *p*_*C*_, are lost at rate 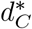 and differentiate at a rate Δ. Upon differentiation, cells undergo *k* rounds of clonal division and a fraction *γ* enter target subpopulation *E*_1_ (with the remainder, 1 *− γ*, entering subpopulation *E*_2_). Cells in the two subpopulation proliferate at rates *p*_1_ and *p*_2_ respectively and are lost at rates *d*_1_ and *d*_2_ respectively. The steady state sizes of the three populations are denoted 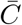, *Ē*_1_ and *Ē*_2_. *U* (*t*) is the availability of label in the body water and *b*_*w*_ the normalisation factor as in equation 2.

### Fitting simulated data (Section *Upstream and downstream compartments*)

When fitting the one compartment model alone to the simulated data (i.e. Section *Confirming that the production by division is estimated accurately using numerical simulation*) then rstan version 2.19.2 was used. Code is written in R version 3.6.3. The prior distributions for *p*_*E*_ and 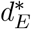 were uniform between 0 and 5*/b*_*w*_, and between 0 and 1 respectively.

When fitting the two compartment precursor/target model we switched to a frequentist approach due to convergence problems with stan. This was implemented via the Pseudo function in the R package FME. In order to ensure fits were comparable, whenever performing a direct comparison with the two compartment model (i.e. Section *Comparing fits using models with and without an upstream compartment*) then the one compartment model was also fitted using FME. The 95% confidence intervals of the parameter estimates were found by bootstrapping the data. That is, 500 bootstrap data sets the size of the original data set were created by randomly sampling with replacement from the original data set. Each bootstrap data set was fitted and the parameters estimated. The 95% CI were taken to be the 0.025 and 0.975 percentiles of the 500 parameter estimates.

### Simulating data for two-compartment explicit heterogeneity model (Section *Kinetic heterogeneity*)

Data were simulated using Eq. 7 with *N* = 2, and all combinations of *p*_1_ = 0.0072, 0.018, 0.036, 0.12, 0.24, 0.36, 0.72; *p*_2_ = 0.0036, 0.0072, 0.0108, 0.0160; *α*_1_ = 0.1, 0.3, 0.5, 0.7, 0.9 where *p*_1_ *> p*_2_. The other parameters were fixed at *δ* = 0.07, *f* = 0.032, *b*_*w*_ = 4.18 and the labelling period is 49 days. Errors on simulated data were lognormally distributed with *σ* = 0.005. Data points were taken every 7 days up to 98 days.

### Fitting data for two-compartment kinetic heterogeneity model (Section *Kinetic heterogeneity*)

The data were fitted using the No U-Turn sampler implemented in the rstan package. Three models were considered: the explicit heterogeneity model (Eq. 7), the implicit heterogeneity model (Eq. 8) and a homogeneous model (i.e. *p* = *d*^*∗*^ in Eq. 8). The body water parameters *f* and *δ*, the amplification factor *b*_*w*_, and the standard deviation for the sampling error, were assumed to be known.

The homogeneous model only has one fitted parameter, *p*, and its prior distribution was uniform between 0 and 1.

The explicit heterogeneity model has 2*N −* 1 fitted parameters, where *N* is the number of compartments. The prior distributions for the proliferation rates *p*_*i*_ are independent; each has a uniform prior of [0, 1]. The prior distribution for *α*_*i*_ is symmetric Dirichlet with concentration parameter *α* = 1. This prior distribution enforces 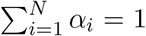. Without loss of generality we order the vector *α*_*i*_ from smallest to largest. For example, for *N* = 2 this is equivalent to a uniform prior for *α*_1_ between 0 and 0.5, and setting *α*_2_ = 1 *− α*_1_. The implicit heterogeneity model has two fitted parameters, *p* and *d*^*∗*^. The prior distribution is independent between parameters, with a uniform prior of [0, 1] for *p* and *d*^*∗*^. The one exception to these priors is in Figure 7B where we explored the impact of prior assumptions. Here the prior distribution for *p* in the homogeneous model is [0, 10]. For the implicit heterogeneity model, priors are [0, 10] for *p* and *d*^*∗*^. For the explicit heterogeneity model with *N* = 2, priors are [0, 10] for *p*_1_ and *p*_2_ (the prior for *α*_1_ is unchanged at [0, 0.5]).

### Model comparison (Section *Kinetic heterogeneity*)

We compare the fits between models with *i* and *j* compartments by computing the point estimate and standard error for Δ*elpd*_*loo*_ = *elpd*_*loo,i*_ *− elpd*_*loo,j*_ using the *loo* package version 2.4.1 in R. *elpd*_*i*_ is the expected log pointwise predictive density for a model with *i* compartments, and *elpd*_*loo,i*_ is its estimate using leave-one-out cross-validation [30]. We define support for model *i* over model *j* when the point estimate for Δ*elpd*_*loo*_ is positive and exceeds its standard error. We define that two models are tied if the standard error for Δ*elpd*_*loo*_ is greater than the magnitude of its point estimate.

### Calculating the discrepancy between label in the blood and in the lymphoid tissue (Section *Spatial distribution of T lymphocytes*)

The discrepancy between label in blood and label in lymphoid tissue was calculated for parameters in the following ranges:

We set 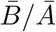, the blood to lymph ratio of interest, to 2*/*98. Bounds on *p* were chosen in line with typical estimates of T cell proliferation in healthy individuals [29]. Our baseline estimate of *f* is calculated from the observation that there are 10^10^ lymphocytes in the blood and that every day 2.5 *×* 10^11^ lymphocytes exit blood for the spleen and 0.3 *×* 10^11^ lymphocytes per day exit the blood for the lymph node; giving an estimate of *f* = 2.8 *×* 10^11^*/*10^10^ = 28*d*^*−*1^. Estimates taken from Kuby Immunology [31] are a similar order of magnitude: there it is reported that it takes 30 mins for a lymphocyte to transit the blood compartment and that 87% of lymphocytes exit for lymphoid tissue giving *f* = 42*d*^*−*1^. These estimates of *f* are then varied over several orders of magnitude (*log*_10_*f* ∈ [−4, 8]) to ensure that results are robust. Given a value of *p* then *d* is constrained by the equilibrium constraint 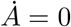 which requires *d* = *p* and so the bounds on *d* are the same as the bounds on *p* but only *p* is independently sampled. Similarly, the equilibrium constraint 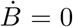 together with the assumption that approximately 2% of lymphocytes are in blood means that 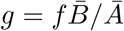.

## Acknowledgements

We are very grateful to Arpit Swain, José Borghans and Rob de Boer for helpful discussions. AY is funded by an Imperial College Research Fellowship. BA is a Wellcome Trust (WT) Investigator (103865Z/14/Z) and is also funded by the Medical Research Council (MRC) (J007439, G1001052), the European Union Seventh Framework Programme (FP7/2007–2013) under grant agreement 317040 (QuanTI), the European Union H2020 programme under grant agreement 764698 (QUANTII) and Leukemia and Lymphoma Research (15012).

## Notes

### Competing Interest Statement

The authors have declared no competing interest.

