## Supplementary Material for "The Impact of Model Assumptions in Interpreting Cell Kinetic Studies"

#### Derivation of approximation used in main text section

To recap the model equations in the main text, the fraction of label in the upstream and downstream compartments are

$$\frac{dF_C}{dt} = p_C b_w U(t) - (d_C^* + \Delta) F_C, \tag{S1a}$$

$$\frac{dF_E}{dt} = (2^k - 1) b_w U(t) \Delta \frac{\bar{C}}{\bar{E}} + \Delta \frac{\bar{C}}{\bar{E}} F_C + p_E b_w U(t) - d_E^* F_E, \tag{S1b}$$

where

$$U(t) = \begin{cases} f[1 - \exp(-\delta t)] & \text{if } t \leq \tau \\ U(\tau) \exp(t - \tau) & \text{otherwise} \end{cases}, \tag{S2}$$

and the fitted model parameters obey

$$\frac{dF_E}{dt} = \hat{p_E} b_w U(t) - \hat{d_E^*} F_E. \tag{S3}$$

Comparing Eqs. S1 and S3, it is unclear how $\hat{p_E}$ and $\hat{d_E}$ relate to $k$, $\Delta$, $p_C$, $d_C$, $p_E$, $d_E$, $\bar{C}/\bar{E}$. This is because $F_C$ is a function of time.

We can analyse this system in two cases:

1. $2^k \gg 1$; and
2. $k = 0$.

# 1 $2^k \gg 1$

We establish an upper bound for $F_C(t)$:

$$F_C(t) = \frac{p_C b_w U(t) - \frac{dF_C}{dt}}{d_C^* + \Delta} \tag{S4a}$$

$$\leq \frac{p_C b_w U(t)}{d_C^* + \Delta} \tag{S4b}$$

$$\leq b_w U(t). \tag{S4c}$$

This result makes sense because the amount of label in cellular DNA cannot exceed the amount of label in body water (multiplied by the normalisation factor, which is the number of hydrogen atoms in deoxyribose available to be replaced by deuterium).

It follows that

$$\frac{dF_E}{dt} \geq (2^k - 1) b_w U(t) \Delta \frac{\bar{C}}{\bar{E}} + p_E b_w U(t) - d_E^* F_E \tag{S5a}$$

$$= \left[ \left( 2^k - 1 \right) \Delta \frac{\bar{C}}{\bar{E}} + p_E \right] b_w U(t) - d_E^* F_E, \tag{S5b}$$

and

$$\frac{dF_E}{dt} \leq (2^k - 1) b_w U(t) \Delta \frac{\bar{C}}{\bar{E}} + b_w U(t) \Delta \frac{\bar{C}}{\bar{E}} + p_E b_w U(t) - d_E^* F_E \tag{S6a}$$

$$= \left( 2^k \Delta \frac{\bar{C}}{\bar{E}} + p_E \right) b_w U(t) - d_E^* F_E, \tag{S6b}$$

As $2^k \gg 1$, the relative difference between $\left(2^k - 1\right) \Delta \frac{\bar{C}}{\bar{E}} + p_E$ and $2^k \Delta \frac{\bar{C}}{\bar{E}} + p_E$ is small. It is then reasonable to conclude, comparing to Eq. S3 that

$$\left( 2^k - 1 \right) \Delta \frac{\bar{C}}{\bar{E}} + p_E \leq \hat{p_E} \leq 2^k \Delta \frac{\bar{C}}{\bar{E}} + p_E, \tag{S7}$$

and

$$\hat{d_E^*} \approx d_E^*. \tag{S8}$$

## 2 $k = 0$

Production by division is defined to be $\Delta(2^k - 1)C/E + p_E$ so in the case $k = 0$ this is equal to $p_E$, the proliferation rate of the target (downstream) population.

The full equation for $F_E$ is given by the second equation of Eq. S1, which in the case $k = 0$ reduces to

$$\frac{dF_E}{dt} = p_E b_w U(t) - (d_E^* F_E - \Delta \frac{\bar{C}}{\bar{E}} F_C) \tag{S9}$$

Given that well-defined numerical values of $b_w$ and $U(t)$ are both entered into this equation then, if we fit the simplified equation, Eq S3, it can be seen that the estimated proliferation rate $\hat{p_E}$ will be a good approximation to $p_E$. It follows that the turnover rate $\Delta C/E + p_E$ will be underestimated by $\hat{p_E}$. It can also be seen that the estimated disappearance rate $\hat{d_E^*}$ will approximate $d_E^* - \Delta \frac{\bar{C}}{\bar{E}} F_C / F_E$, since the last term is always positive this will lead to an underestimate of $d_E^*$.

29 # Supplementary Figures

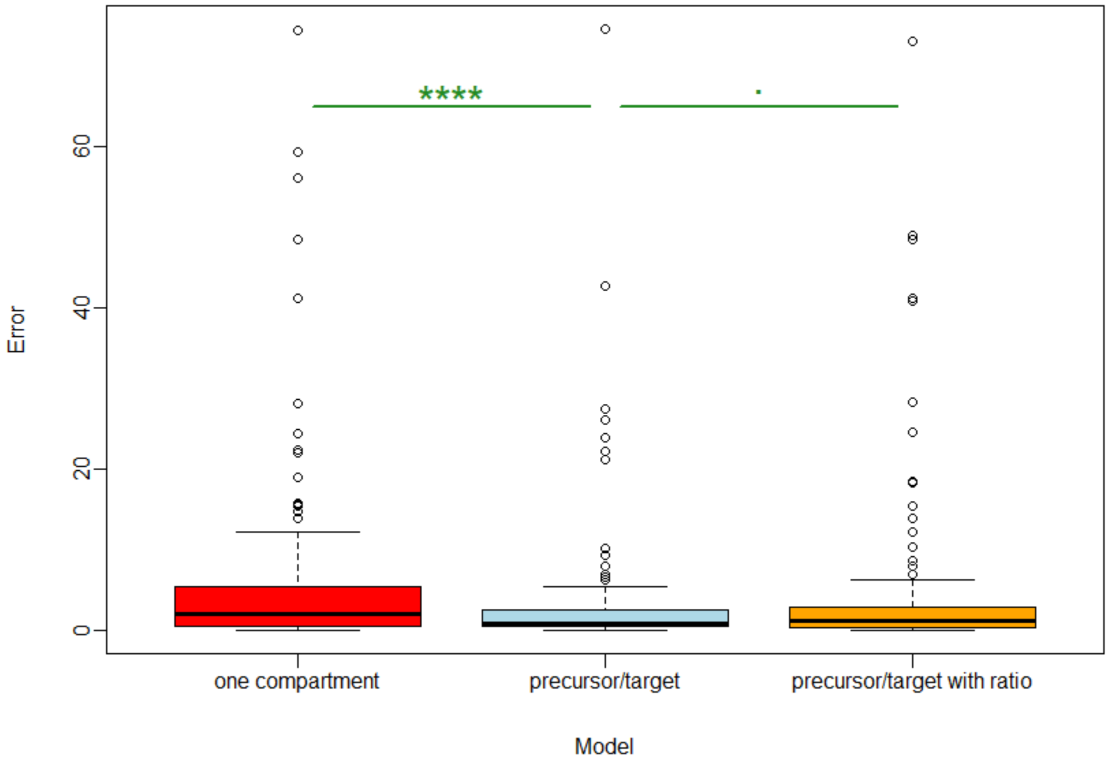


**Supp. Figure 1. Error in estimates of the proliferation rate.** This shows the same data as in Figure 4A but without the y axis truncation.

A

B

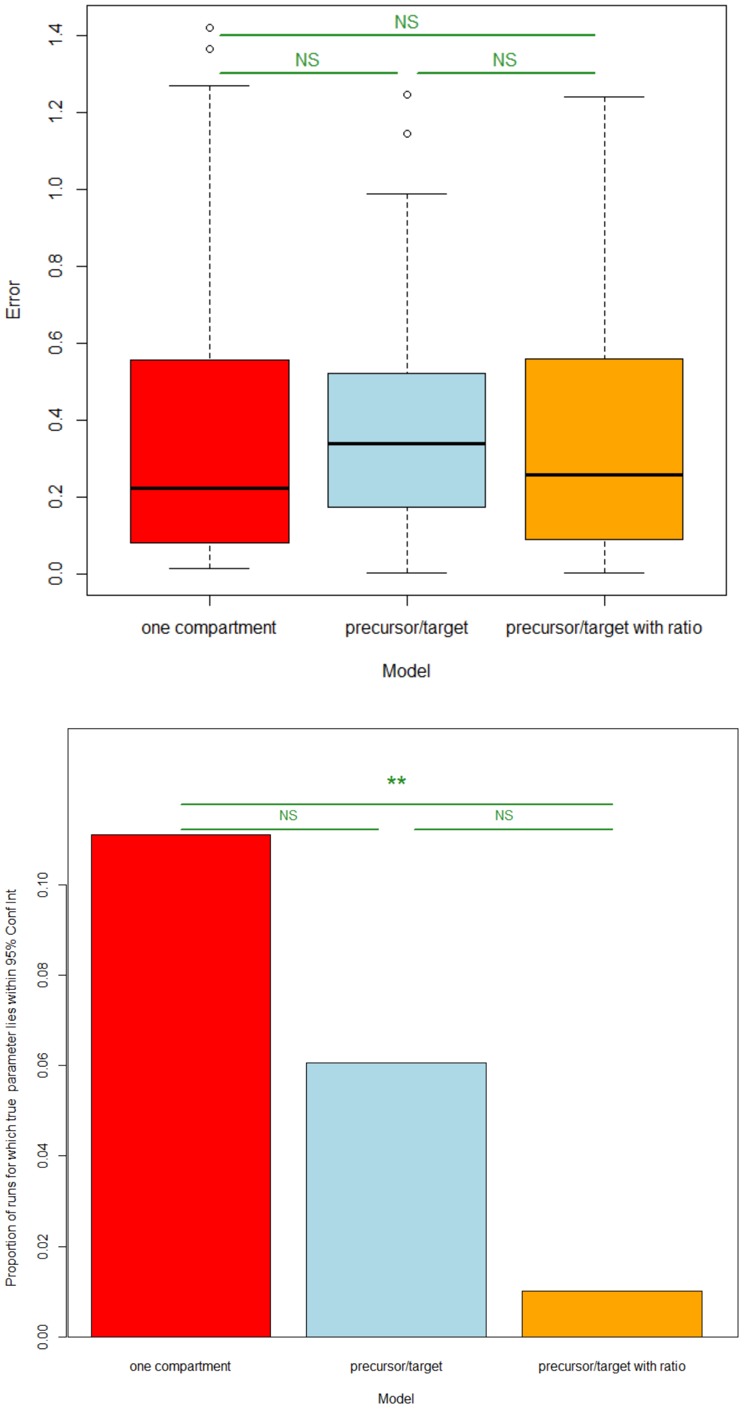


**Supp. Figure 2. Error in estimates of the proliferation rate for realistic data but with a high proliferation rate. A** Discrepancy between point estimate of the proliferation rate and the true value (expressed as a fraction of the true value) for the one compartment model (red), precursor/target model (blue) and precursor/target with ratio model (orange). On increasing the values of $p_E$ used to generate the labelling data two effects were noticed compared to the results depicted in Figure 4: first, errors were now much lower, second, there was no longer any significant difference in the size of the errors associated with each of the three models. **B** 95% confidence intervals (CI) were estimated by bootstrapping the data (Methods) and the fraction of runs where the true value lay within the CI was reported. Colours as for A. Compared to the corresponding figure for realistic data but with a lower value of $p_E$ (Figure 4) the proportion of runs where the estimate fell within the CI was still very low but the pattern across the models was quite distinct, for this set of parameters the one compartment model outperformed the other two models. Significance codes: NS $P > 0.05$, * $0.01 < P \leq 0.05$, ** $0.001 < P \leq 0.01$, ** *$0.0001 < P \leq 0.001$, **** $0.00001 < P \leq 0.0001$

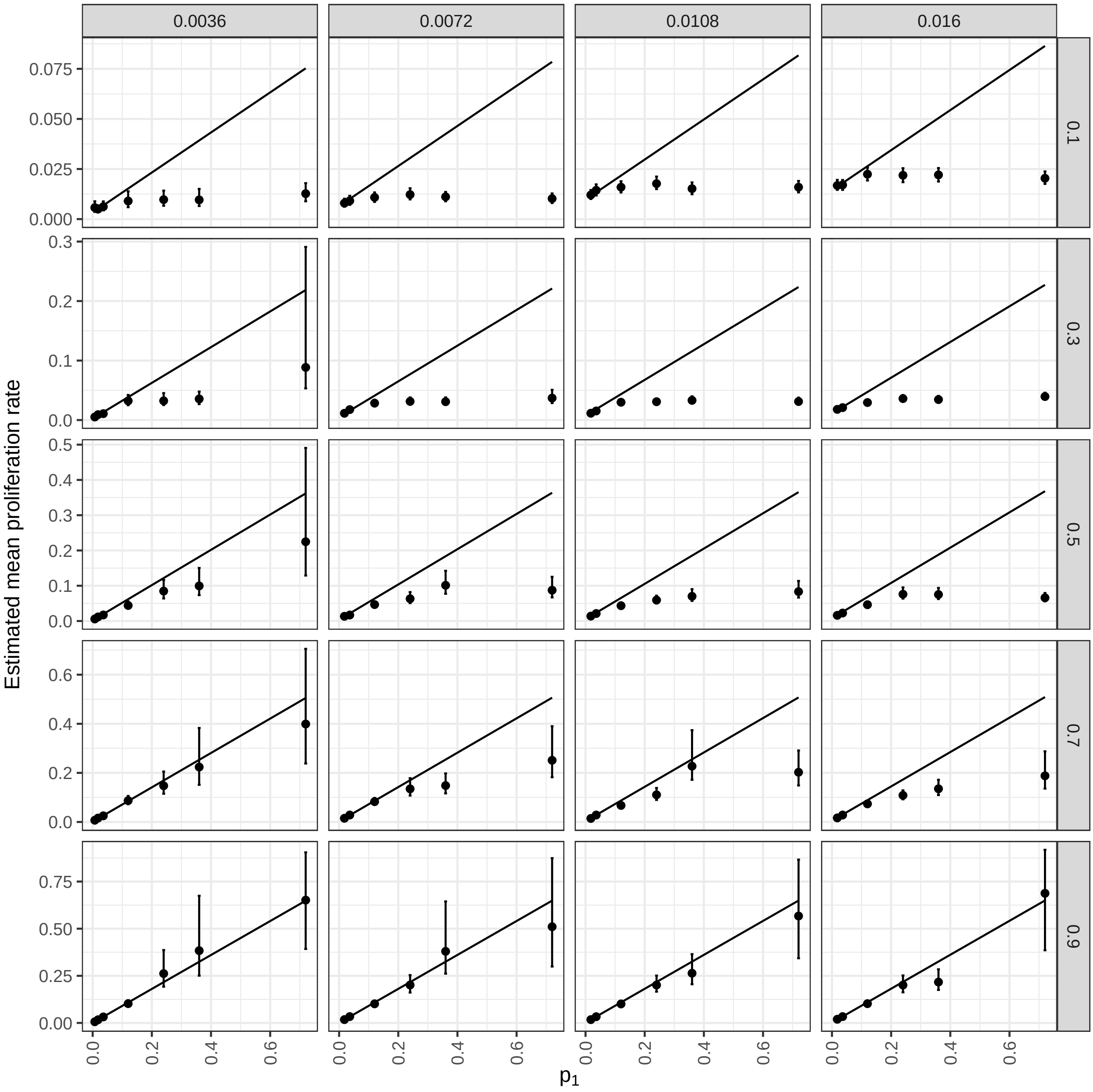


**Supp. Figure 3. Median and 95% credible intervals for the mean proliferation rate when fitting the implicit model to data generated using the two-compartment explicit model.** The value of $p_1$ used to simulate the data is shown on the x-axis. Within each plot, the values of $p_2$ and $\alpha_1$ used to simulate the data are held constant at the values on the top and right respectively.

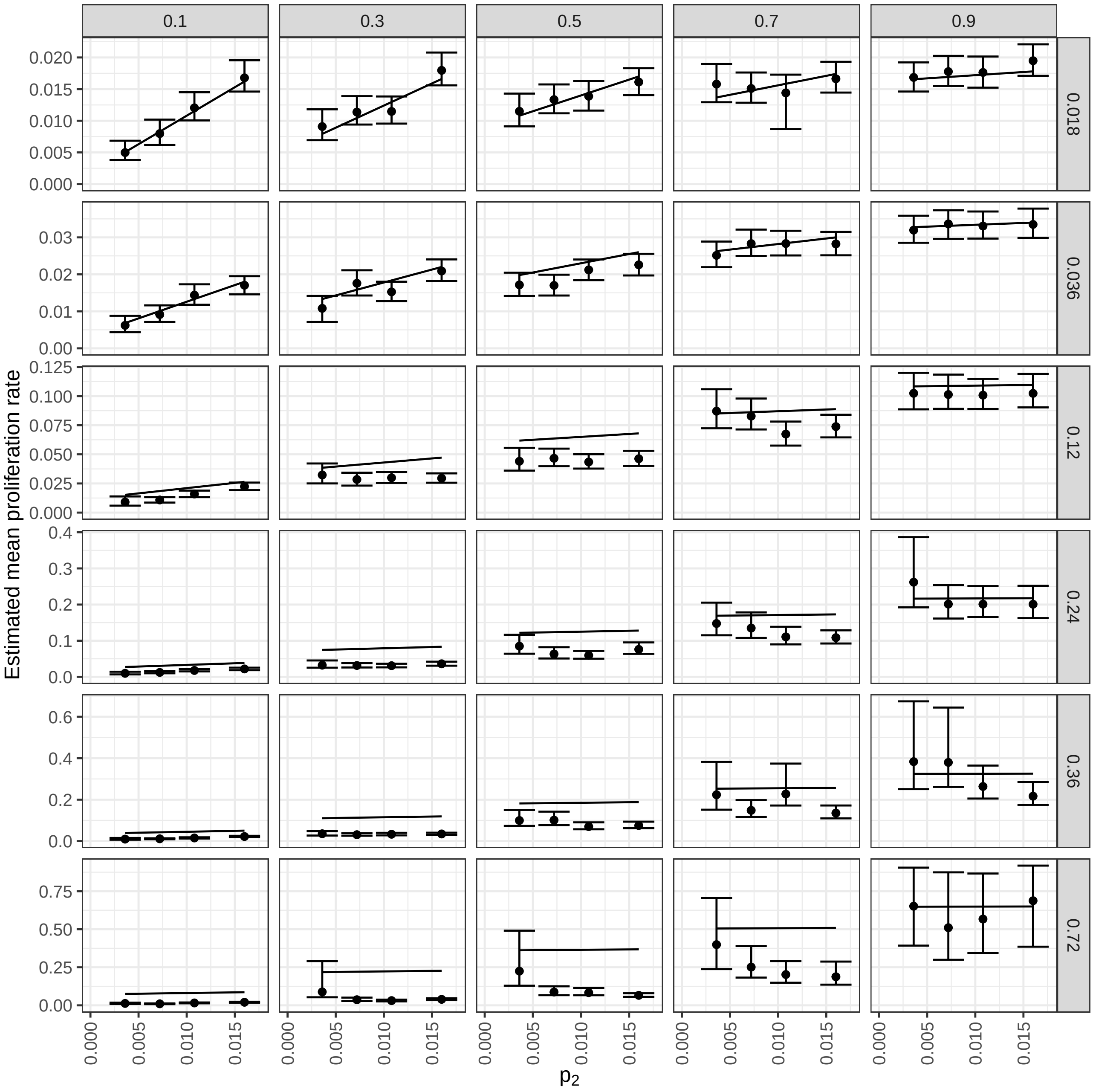


**Supp. Figure 4. Median and 95% credible intervals for the mean proliferation rate when fitting the implicit model to data generated using the two-compartment explicit model.** The value of $\alpha_1$ used to simulate the data is shown on the x-axis. Within each plot, the values of $\alpha_1$ and $p_1$ used to simulate the data are held constant at the values on the top and right respectively.

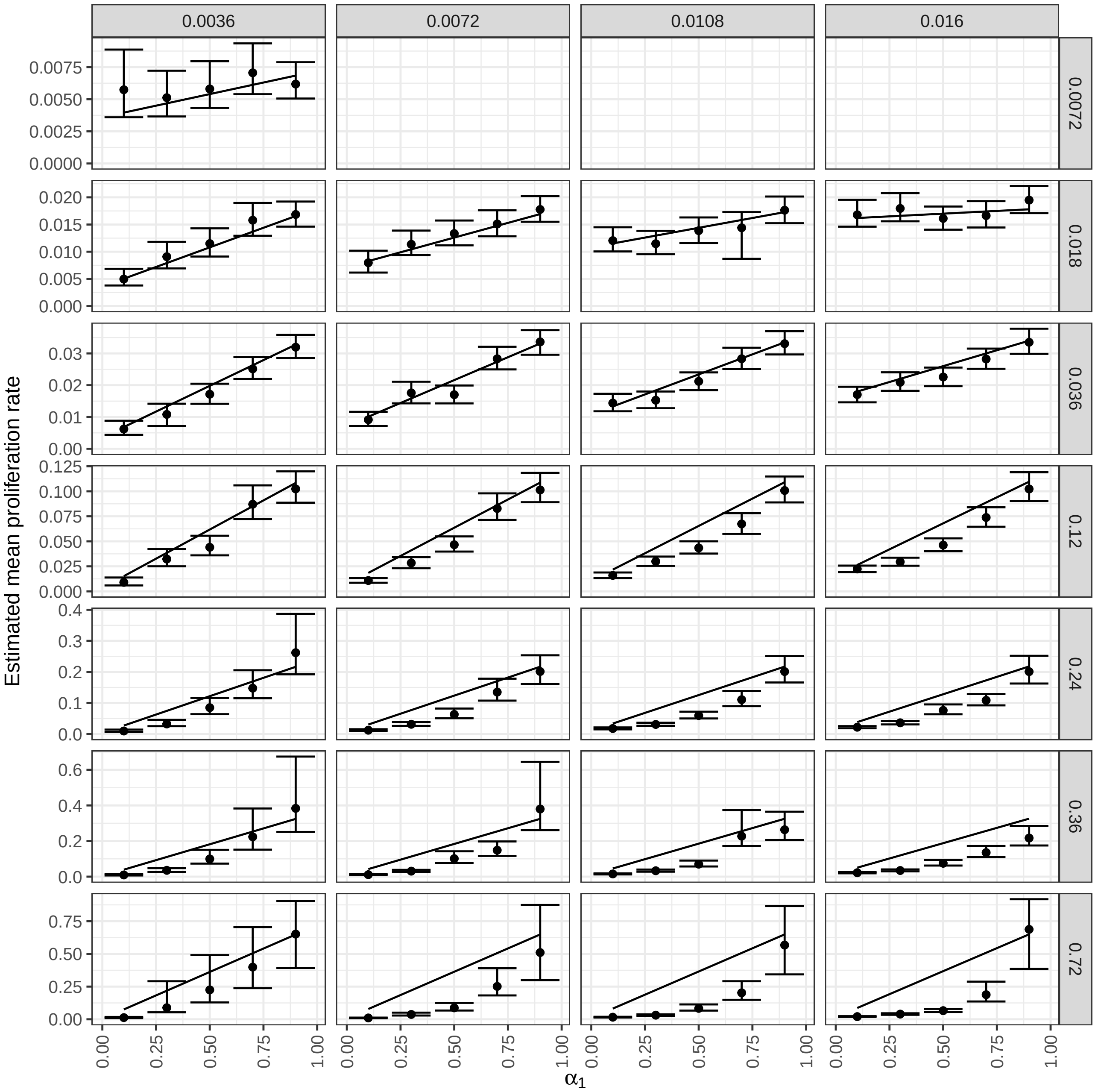


**Supp. Figure 5. Median and 95% credible intervals for the mean proliferation rate when fitting the implicit model to data generated using the two-compartment explicit model.** The value of $\alpha_1$ used to simulate the data is shown on the x-axis. Within each plot, the values of $p_2$ and $p_1$ used to simulate the data are held constant at the values on the top and right respectively.

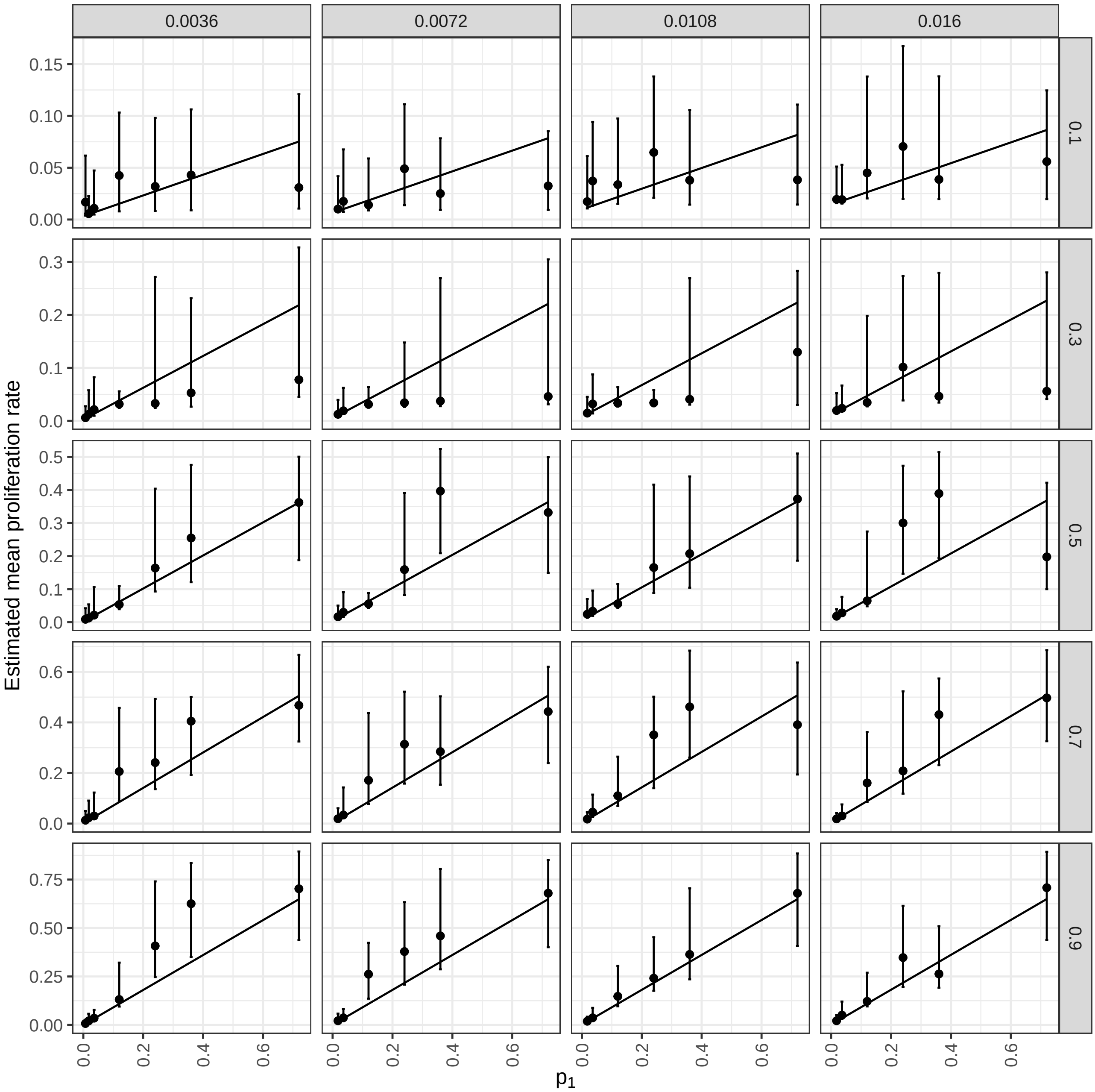


**Supp. Figure 6. Median and 95% credible intervals for the mean proliferation rate when fitting the two-compartment explicit model to data generated using the same model.** The value of $p_1$ used to simulate the data is shown on the x-axis. Within each plot, the values of $p_2$ and $\alpha_1$ used to simulate the data are held constant at the values on the top and right respectively.

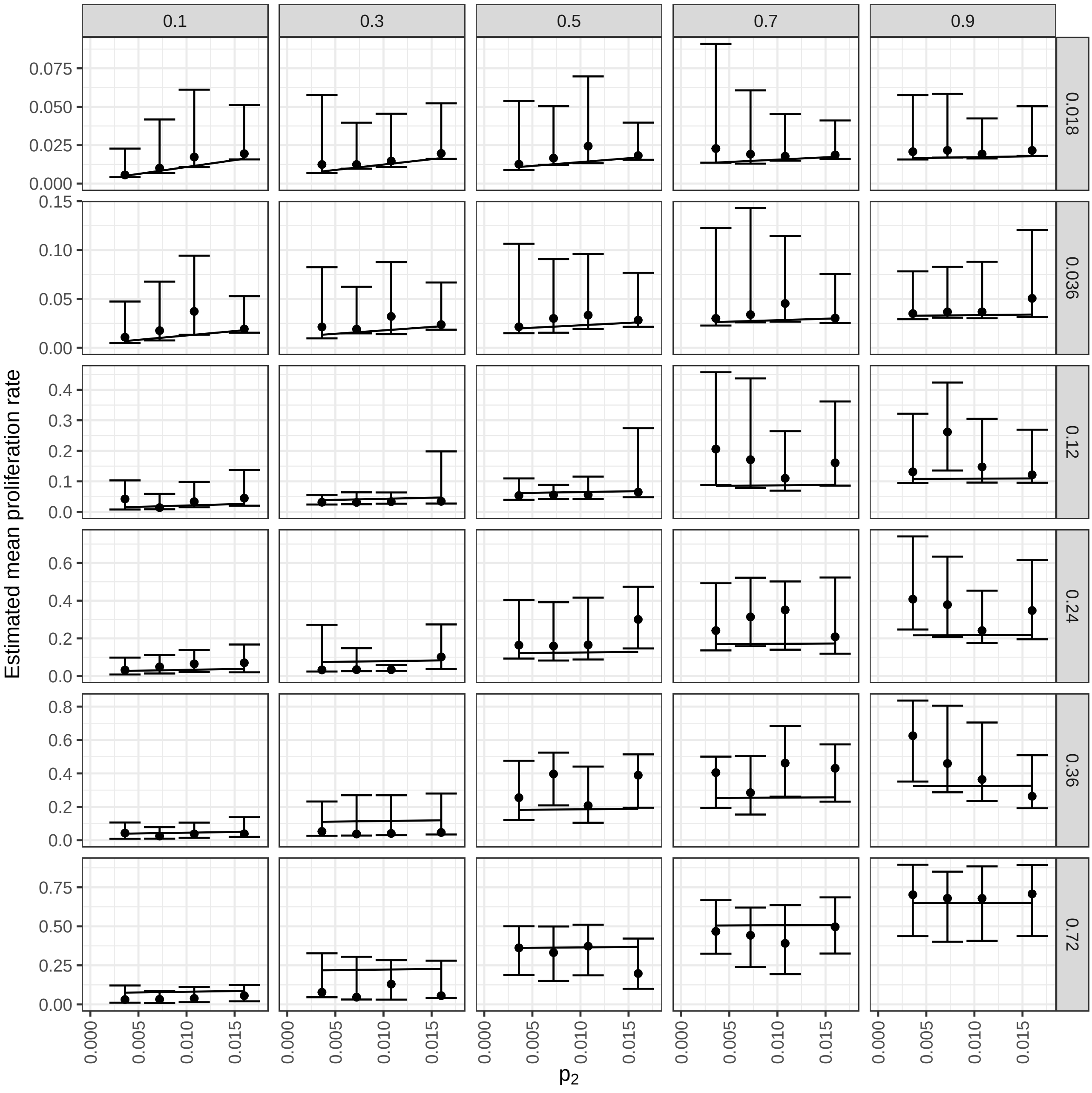


**Supp. Figure 7. Median and 95% credible intervals for the mean proliferation rate when fitting the two-compartment explicit model to data generated using the same model.** Median and 95% credible intervals for the mean proliferation rate when fitting the the two-compartment explicit model to data generated using the same model. The value of $\alpha_1$ used to simulate the data is shown on the x-axis. Within each plot, the values of $\alpha_1$ and $p_1$ used to simulate the data are held constant at the values on the top and right respectively.

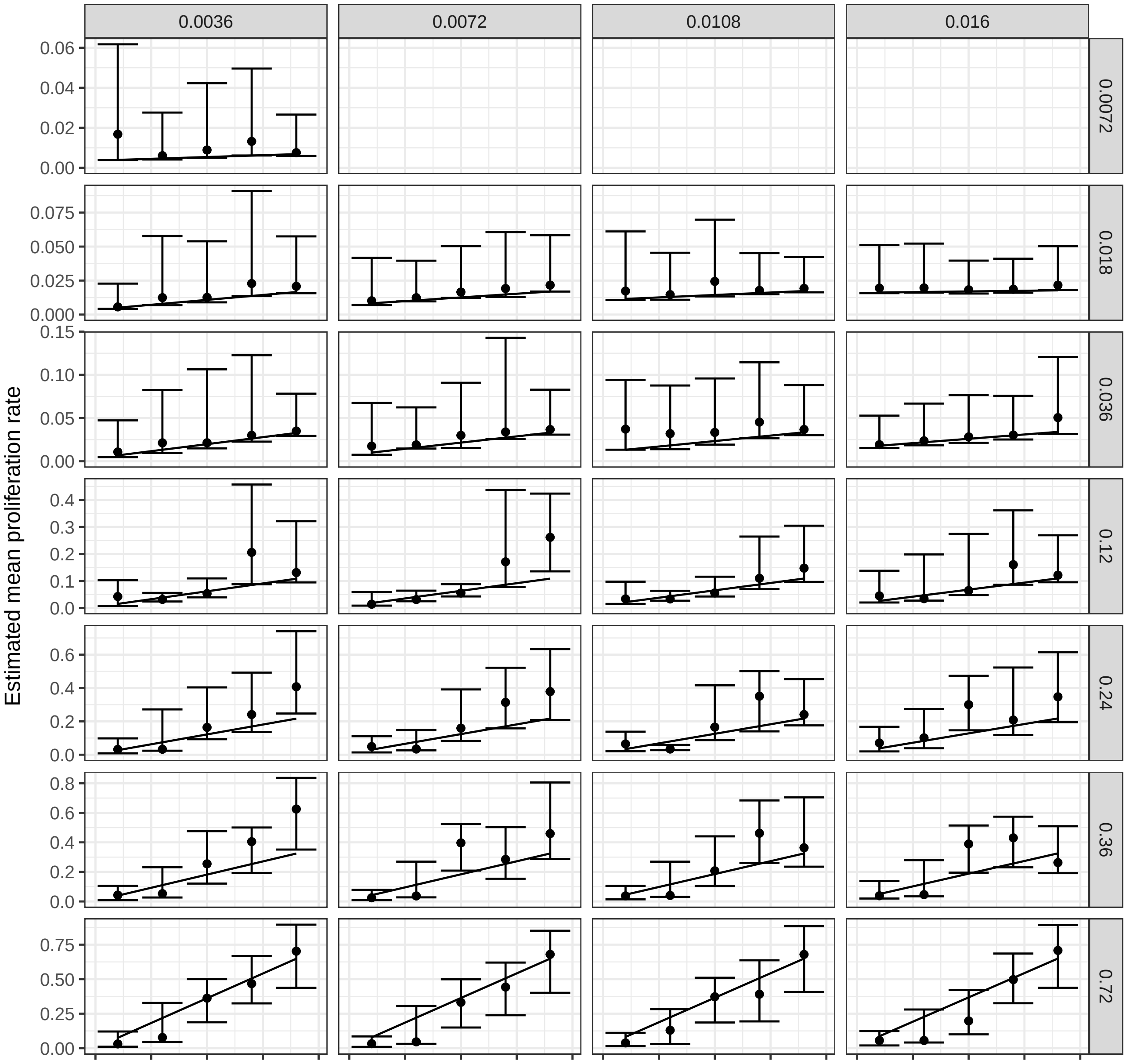


**Supp. Figure 8. Median and 95% credible intervals for the mean proliferation rate when fitting the two-compartment explicit model to data generated using the same model.** Median and 95% credible intervals for the mean proliferation rate when fitting the the two-compartment explicit model to data generated using the same model. The value of $\alpha_1$ used to simulate the data is shown on the x-axis. Within each plot, the values of $p_2$ and $p_1$ used to simulate the data are held constant at the values on the top and right respectively.

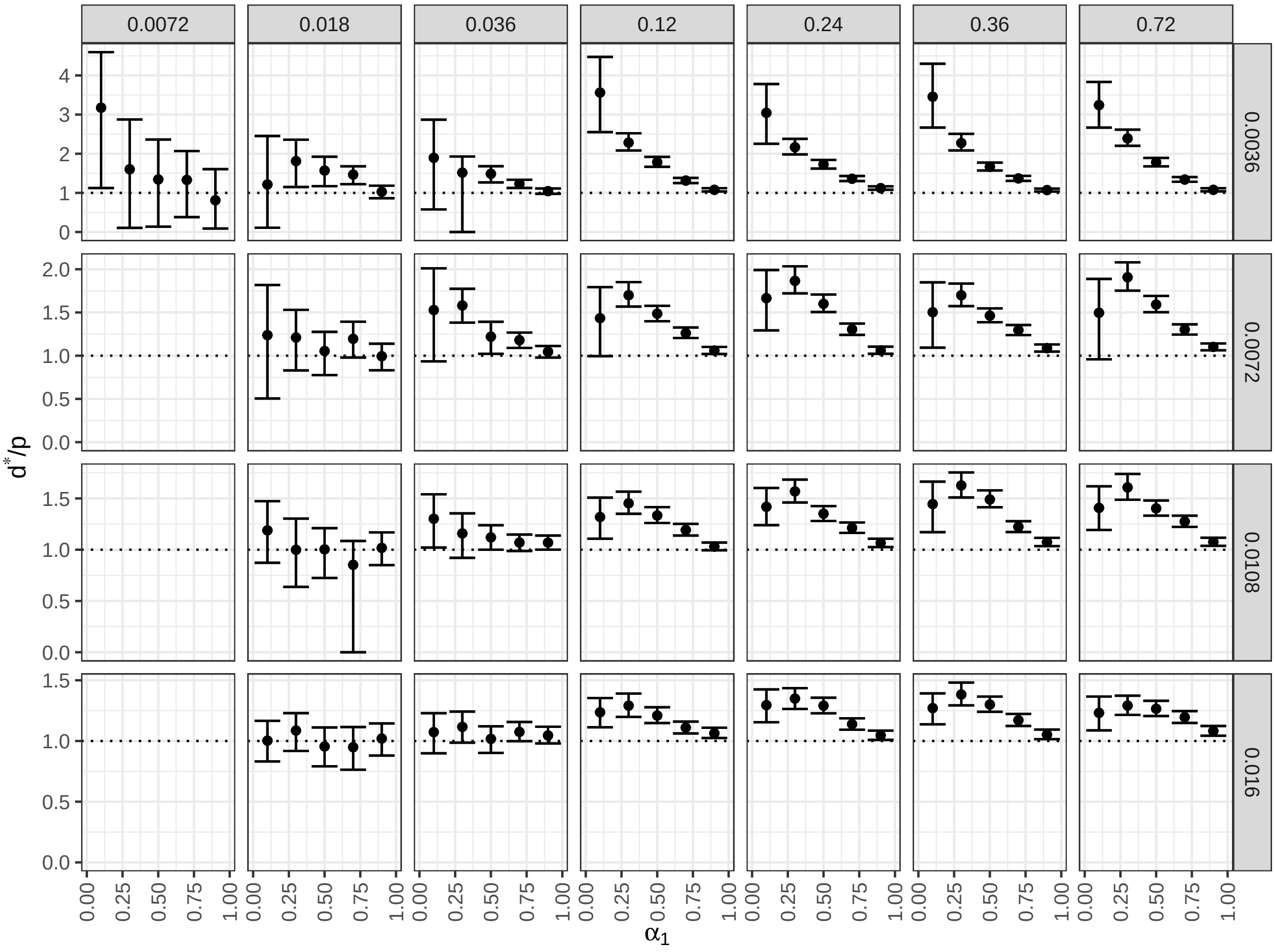


**Supp. Figure 9. The 95% credible intervals for $d^*/p$ when fitting the implicit kinetic heterogeneity model to data generated using the two-compartment explicit kinetic heterogeneity model.** If the 95% credible interval excludes 1, then there is significant evidence for kinetic heterogeneity. The data were generated using different values of $\alpha_1$ (x-axis), $p_1$ (label to right of plots) and $p_2$ (label above plots). There is significant evidence for kinetic heterogeneity for high values of $p_1$, with more evidence at lower values of $\alpha_1$.

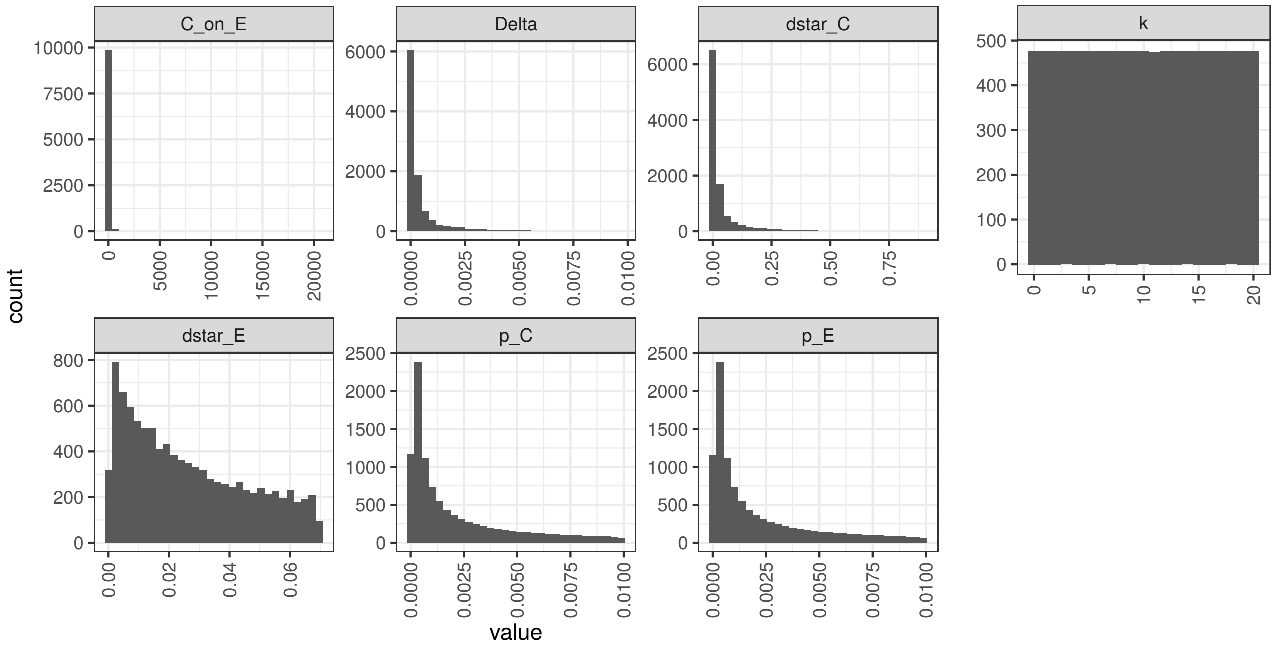


**Supp. Figure 10. Distribution from which simulation parameters are sampled for Section .**

### Supplementary Tables

| Parameter | Value |
|---|---|
| $p_C$ | $4.50 \times 10^{-4}$ |
| $p_E$ | $2.39 \times 10^{-3}$ |
| $k$ | $0, 10$ |
| $\Delta$ | $1.50 \times 10^{-4}$ |
| $d_C^*$ | $3 \times 10^{-4}$ |
| $d_E^*$ | $7.75 \times 10^{-3}$ |
| $\bar{C}/\bar{E}$ | 32.16 |
| $f$ | 0.032 |
| $\delta$ | 0.070 |
| $b_w$ | 5.82 |

**Supp. Table 1.** Parameter values used to simulate the data underlying Figure 2. All rates have units day$^{-1}$.

| Parameter | A, D | B, E | C, F |
|---|---|---|---|
| $p_1$ | varied | 0.018 | 0.72 |
| $p_2$ | 0.016 | varied | 0.016 |
| $\alpha_1$ | 0.7 | 0.1 | varied |
| $f$ | 0.032 | | |
| $\delta$ | 0.064 | | |
| $b_w$ | 4.18 | | |

**Supp. Table 2.** Parameter values used to simulate the data underlying Figure 6. All rates have units day$^{-1}$.
